# Distributed neural dynamics underlie the shift from movement preparation to execution

**DOI:** 10.64898/2025.12.09.693175

**Authors:** Ziwei Yin, Jian K. Liu, Katja Kornysheva

## Abstract

Understanding how neural population activity supports the transition from movement preparation to execution remains a central challenge in systems neuroscience. Invasive recordings in non-human primates indicate that motor preparation and execution occupy orthogonal subspaces within the primary motor cortex (M1), yet how these neural dynamics evolve across broader networks and support skilled, memory-guided action remains poorly understood. We used source-resolved magnetoencephalography (MEG) during a memory-guided finger-sequence task to track population dynamics across a cortico–hippocampal network. Manifold trajectories and decoding revealed that preparation and execution of movement sequences occupy distinct, yet not fully orthogonal, neural subspaces – a separation that generalised across all regions but was most pronounced in motor regions. Crucially, we identified a hierarchical spatiotemporal transition across the brain: the state shift originated in the hippocampus via theta-band modulations and rippled through the pre-SMA, SMA, and lateral premotor areas before culminating in M1 via broadband activity. This progression follows the functional hierarchy of descending motor pathways, with M1 shifting last, 160 ms before the first button press. Although sequence identity could be decoded from all areas during execution, only the hippocampus and M1 maintained predictive representations of upcoming sequences during preparation. These findings demonstrate that the initiation of complex motor sequences relies on a well-timed integration of hippocampal memory systems with primary motor output, rather than being a process localised to the motor cortices alone.

## Introduction

Coordinating skilled actions requires the nervous system to prepare upcoming movements while preventing their premature release (Churchland, 2024). In non-human primates, invasive recordings in the primary motor cortex (M1) and adjacent areas have shown that preparation and execution are implemented through distinct population-level neural states that unfold within separate low-dimensional subspaces of the same motor population (Churchland et al., 2012; Elsayed et al., 2016; Kaufman et al., 2014). Preparatory activity evolves in an “output-null” space, enabling the retrieval of motor plans without immediate motor output, while execution occupies an “output-potent” space defined by characteristic rotational dynamics (Churchland & Shenoy, 2024; Churchland et al., 2012; Kao et al., 2021; Kaufman et al., 2014). These findings support a dynamical-systems view in which actions emerge from evolving population trajectories, i.e., dynamic response patterns rather than static tuning of individual neurons, activated above or below a threshold, to upcoming movement features (Shenoy et al., 2013).

In humans, neuroimaging can reveal large-scale activity pattern transitions from preparation to execution across the brain. For example, fMRI results suggest that the control of sequence features undergoes a state shift from high-level feature separation during planning to their integration during execution across a network of motor-related cortical and subcortical areas (Yewbrey & Kornysheva, 2024; Yewbrey et al., 2023). However, the temporal dynamics and geometry of this transition remain unknown. MEG/EEG provide non-invasive access to brain dynamics, including well-established oscillatory markers of movement preparation, such as beta and alpha (mu)-band desynchronisation before and during movement linked to cortical excitability, followed by their post-movement rebound linked to inhibition (Barone & Rossiter, 2021; Engel & Fries, 2010; Jurkiewicz et al., 2006; Muthukumaraswamy et al., 2004), as well as theta-band activity associated with cognitive control and memory-dependent planning (Dash et al., 2025; Hannah et al., 2018). However, these oscillatory dynamics alone do not reveal whether preparation and execution patterns occupy distinct neural manifolds, nor how the transition between them unfolds across distributed brain regions.

Skilled actions typically require the integration of hierarchical information spanning movement sequence structure, chunk organisation and individual movements (Dash et al., 2025; Kikumoto & Mayr, 2018). Prior work shows that these levels are not represented uniformly across the motor system: the primary motor cortex primarily encodes upcoming single movements (Ariani et al., 2022; Dekleva et al., 2018; Zimnik & Churchland, 2021), whereas lateral (PMd, PMv) and medial premotor regions (SMA, pre-SMA) contain sequence-selective activity (Shima & Tanji, 2000; Yewbrey et al., 2023; Yokoi & Diedrichsen, 2019). Further, there is growing evidence of the hippocampus supporting retrieval and control of sequential order (Dolfen et al., 2024; Reverberi et al., 2026; Yewbrey & Kornysheva, 2024), with hippocampal–cortical replay predicting short-term performance gains (Buch et al., 2021). Collectively, this broadly distributed architecture indicates that motor sequence control engages both cognitive and motor neural systems spanning a wide anatomical gradient of proximity to the motor periphery. This gradient is anatomically anchored by the density of direct corticospinal projections to lower cervical spinal segments (C7–T1) supplying hand motor neurons and reciprocal cortico-cortical connectivity with M1 digit representations (Dum & Strick, 2005). However, invasive studies have largely focused on M1 and premotor regions, leaving it unclear how these and other motor sequence-relevant regions coordinate the transition from preparation to execution. Whether the preparatory–execution manifolds observed in M1 extend to more complex action sequences and to motor cortical and hippocampal networks as suggested by fMRI in humans (Yewbrey & Kornysheva, 2024; Yewbrey et al., 2023), and how their dynamics unfold in time, remains unknown.

Here, we examined source-resolved MEG obtained during a memory-guided finger-sequence task to probe the preparatory–execution state transition non-invasively and assess its organisation across motor, premotor and hippocampal regions. We further asked whether oscillatory MEG dynamics track the transition and support the decoding of upcoming sequence identity. Our results reveal a global shift across distinct manifolds toward execution-related patterns prior to movement onset, with evidence of hierarchical timing across premotor and primary motor regions, and sequence-specific information detectable in hippocampal and primary motor areas even before movement execution. These results demonstrate that non-invasive MEG can resolve continuous, hierarchical state transitions and reveal distributed population dynamics underlying sequential action in the human brain.

## Materials and Methods

### Memory-guided finger sequence task

This study reanalysed MEG data from Kornysheva et al. (2019). Participants were trained over two days to associate four abstract visual cues, drawn from a pool of 16 fractal images, with four five-finger tapping sequences. Each sequence combined one of two finger orders (F1, F2) with one of two temporal interval orders (T1, T2; 550, 650, 800, 983, and 1300 ms), yielding a 2 × 2 factorial design. For each participant, the identity of the first finger press remained constant across all four sequences. The finger-order and temporal-interval sequences were independently generated for each participant; consequently, nominal sequence identities were not matched across participants. Each trial began with a visual sequence cue, followed by a variable delay of 1.8, 2.0, or 2.2 s, after which participants initiated the memorised sequence in response to the Go Cue.

On day 1, training comprised seven instructed blocks (168 trials), seven mixed blocks, and seven memory-based blocks (168 trials). The mixed blocks contained 56 instructed and 112 memory trials in a blocked 1:2 ratio. In instructed trials, a circle appeared on the target finger at the target time after the Go Cue, and participants synchronised their responses to it. On day 2, participants completed seven mixed blocks, as on day 1, followed by 12 memory-based blocks (288 trials). Additional testing was conducted before and after training on days 1 and 2, respectively, using instructed trials with both trained and untrained sequences. Untrained sequences used novel finger and interval orders.

After each trial, participants received feedback showing their cumulative score and whether they had used the correct finger at the correct time. They earned two points for a correct sequence performed within 30% of the target timing, one point for a deviation below 60%, and zero points otherwise. During the first two training days, auditory feedback indicated whether participants had earned 0–2 points. No auditory feedback was provided during the day 3 MEG session.

The MEG session began with two mixed blocks (16 instructed and 32 memory trials) to refresh participants’ memory of the sequences and familiarise them with the MEG environment. This was followed by 10 blocks of concurrent MEG and EMG recording, comprising 240 memory-based trials distributed equally across the four trained sequences. Trials were arranged in groups of three consecutive repetitions, with group order randomised throughout the session. Throughout, participants were instructed to execute the sequences as accurately as possible entirely from memory without relying on visual prompts for individual presses.

MEG data were continuously recorded at 1,200 Hz using a 275-channel axial-gradiometer system (CTF Omega, VSM MedTech) in a magnetically shielded room. Following the original study, sensor-level data were preprocessed using independent component analysis (ICA) for artefact removal and downsampled to 1,000 Hz. Trial epochs extended from 2.8 s before to 12 s after sequence-cue onset.

For the pre- and post-test instructed trials, a correct trial was defined by the required finger-press sequence and the timing of each response relative to its corresponding finger cue. For memory-based trials, correctness was defined by the required finger-press sequence and reproduction of the target interval structure, quantified as the mean absolute deviation from the target intervals and expressed as a percentage. Only trials with temporal deviations of less than 3 standard deviations were considered accurate and included in further analyses.

Of the 16 healthy right-handed participants included in the original study (Kornysheva et al., 2019), 14 were retained for the present analysis. One participant was excluded because of excessive motion artefacts, and another because of an imbalance in sequence accuracy across finger sequences. While the number of participants is small for an MEG study, the study utilises within-subject dense sampling (60 trials per sequence per participant = 240 trials/session), maximising statistical power at the individual-participant level prior to group inference.

### Statistical analysis

Statistical analyses were performed using non-parametric methods. Group-level omnibus effects were assessed using the Friedman test. Pairwise comparisons were conducted using Wilcoxon signed-rank tests. One-sample comparisons against reference values were performed using one-sample Wilcoxon signed-rank tests. The family-wise error rate was controlled using the Bonferroni correction for fewer than 10 comparisons and the Benjamini–Hochberg false discovery rate (FDR-BH) procedure for 10 or more comparisons. For comparisons between V1 and the other areas, pairwise tests were first performed across all area pairs (including pairs of motor regions) and then corrected using FDR-BH; only significant pairs involving V1 are shown. Time-resolved statistics were evaluated using one-sided, paired cluster-based permutation test (10000 repeats) with a cluster-forming threshold of *p* < .01; clusters with *p* < .05 after false discovery rate correction were considered significant.

#### Linear mixed-effects models

To test whether neural measures followed a motor hierarchy in sequence control, regions were assigned ordinal ranks based on their anatomical corticospinal projections to the lower cervical spinal cord (C7–T1) and cortico-cortical connectivity with M1(Dum & Strick, 2005). We then tested whether multiple metrics followed this proposed regional hierarchy. The hierarchy was ordered as follows: hippocampus (rank 1), pre-SMA (rank 2), SMA (rank 3), PMv (rank 4), PMd (rank 5), and M1 (rank 6).

For each outcome measure and analysis phase, we fitted a linear mixed-effects model of the form

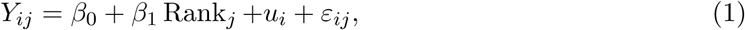

where *Y_ij_* denotes the outcome for participant *i* in region *j*, *β*_0_ is the intercept, *β*_1_ is the linear effect of hierarchical rank, *u_i_* is a participant-specific random intercept, and *ε_ij_* is the residual error. Each participant contributed one value per included region, such as mean decoding accuracy or transition time.

The slope *β*_1_ tested whether the outcome changed systematically along the hierarchy. For transition times, a negative slope indicated progressively earlier transitions toward M1, whereas a positive slope indicated the opposite pattern.

Models were fitted in Python using the statsmodels MixedLM implementation with the default restricted maximum-likelihood estimator. Two-tailed tests of *H*_0_ : *β*_1_ = 0 were performed, with statistical significance defined as *α* = 0.05.

### MEG source reconstruction

Sensor data were co-registered to a standard template MRI in FieldTrip (http://www.fieldtriptoolbox.org/; v20260214) using fiducial-based alignment of the nasion and the left and right preauricular points. A single-shell volume-conduction model and a regular volumetric source grid with 1-cm spacing were used to compute the leadfield matrix.

Prior to source reconstruction, sensor data were baseline-demeaned using the 1-s interval preceding sequence-cue onset. Source activity was estimated using a linearly constrained minimum variance (LCMV) beamformer. For each participant, a common spatial filter was computed at every source-grid location using the covariance matrix from the entire 14.8-s trial, averaged across all correctly performed trials. This covariance estimate, together with the forward model, was used to derive a single spatial filter per source-grid location. The same filters were then applied across all trials and experimental conditions to ensure consistent source reconstruction.

Motor regions (M1, PMd, PMv, SMA, pre-SMA) were defined using the Human Motor Template (HMAT) atlas (Mayka et al., 2006), and the hippocampus using NeuroVault sub-cortical masks (Gorgolewski et al., 2015). The V1 control region was defined using the hOc1/Area 17 probabilistic cytoarchitectonic map from the Jülich Brain Atlas (Amunts et al., 2002, 2020). Grid points belonging to each ROI were identified by anatomical labels and aggregated by region. Source time series were extracted at 1,000 Hz. The source channels were sorted and filtered using the anatomical coordinates to ensure consistent and comparable source channels across participants.

### Time–frequency analysis and cluster-based statistical testing

Time–frequency representations (TFRs) were computed from the source-localised data (1,000 Hz) using the hanning method. Data were aligned to the Go Cue without truncation and mirror-padded to 40 s to prevent edge artefacts. Spectral power was estimated from 0.5 to 80 Hz in 0.5 Hz steps and 50 ms timesteps. A Hanning taper with a window length of six cycles per frequency was applied.

For visualisation, TFRs were dB-normalised relative to each participant’s 1-s baseline preceding the earliest sequence cue and then grand-averaged across participants. Group-level baseline demeaning was performed to centre the baseline grand average at zero.

Statistical evaluation of task-related power changes was performed using a cluster-based permutation test (FieldTrip function ft_freqstatistics). The 1–s baseline period was repeated to match the task window length. Significance was assessed using a Monte Carlo method with dependent-samples t-statistics (cfg.method = ‘montecarlo’, cfg.statistic = ‘depsamplesT’, cfg.correctm = ‘cluster’). The cluster-forming threshold was set at clusteralpha = .01, and post-permutation clusters were considered significant at alpha = .05.

### Linear discriminant analysis of phase prediction

Linear Discriminant Analysis (LDA) was applied to source-reconstructed time series, downsampled to 200 Hz. For each participant, the classifier was trained to discriminate between eight conditions defined by a 2 *×* 4 factorial design: task phase (preparation vs execution) crossed with sequence identity (sequences 1–4). This yielded the following labels: prep*×*seq1, exec*×*seq1, …, prep*×*seq4, exec*×*seq4. The preparation period was defined from sequence cue onset to the variable Go Cue; execution spanned from the first to the final finger press, defined based on trial-wise variable timing.

Decoding results were averaged across 1,000 bootstraps. For each iteration, 40 correctly performed trials were sampled per sequence, except for P02, for whom 15 trials were sampled; these were split into training and test sets (70*/*30 ratio), with stratification to maintain sequence balance. Samples spanning 1 s were randomly extracted from either the preparation or execution period of each training trial and labelled by phase–sequence identity. The trained classifier was then applied continuously to the full test-trial time course to obtain decoding probability trajectories.

Classification accuracy values represent the time-resolved accuracy of assigning the correct phase–sequence label, calculated exclusively from the test trials of the matching sequence. Chance-level baselines were generated by repeating this procedure with randomly permuted class labels while keeping all other processing steps constant. For visualisation, accuracy curves were smoothed using a Gaussian kernel (*σ* = 25 ms) and averaged across iterations within each participant. Group means and standard errors of the mean (SEM) were then calculated across participants.

To quantify the timing of phase transitions, decoding curves were aligned with task events to account for the variable Go Cue and press timing. Go Cue-aligned prediction curves were used to measure the offset of preparation patterns, and first-press-aligned curves to measure the onset of execution patterns. Cluster-based permutation significance was computed per participant using 1,000 bootstraps of observed trials and permutation probability (clusteralpha = .01, final cluster alpha = .05, Bonferroni-correction over 2 phases). Significant clusters shorter than 10 ms were discarded, and clusters separated by less than 10 ms were merged before extracting timing estimates.

### Regression of canonical frequency power onto LDA-derived state estimates

To relate frequency-specific neural activity to the emergence of preparation and execution patterns, LDA predictions were regressed against canonical frequency band power. Time–frequency representations (TFRs) were computed for each participant using the same settings as above, but at high temporal resolution (200 Hz) to match both the sampling rate and temporal structure of the LDA output. Spectral power was extracted for five canonical frequency bands: delta (1 *−* 4*Hz*), theta (4 *−* 8*Hz*), alpha (8 *−* 13*Hz*), beta (13 *−* 30*Hz*), and gamma (30 *−* 80*Hz*). TFRs were z-scored per trial and then averaged across trials within each sequence and frequency band.

To examine the relationship between spectral dynamics and execution-related decoding, LDA predictions were z-scored and then averaged across 1,000 bootstrap repeats per sequence. Time-resolved regressions were computed using a sliding-window approach with a 200-ms window and a 10-ms step. For each window, a predictor matrix *X ∈* R^160×5^ (4 sequences *×* 40 timepoints *×* 5 bands) was regressed against the corresponding LDA predictions *y ∈* R^160^ (4 sequences *×* 40 timepoints) using ridge regression with a regularisation parameter of *α* = 1.

Statistical significance was assessed using the cluster-based permutation test against the permuted regression curve (cluster-forming alpha = .01; corrected alpha = .05, Bonferroni-corrected over 5 frequencies). The permutation regression curve was generated by circularly shifting the predictor time series with a minimum shift of 10% of the trial time length across 1000 repeats.

A similar analysis was performed to assess the relationship between spectral dynamics and sequence identity predictions. In this case, the LDA classifier already yielded time-resolved sequence predictions using the same sliding-window configuration (200 ms window, 10 ms step). The spectral predictors were temporally aligned to the corresponding LDA output for each window. The predictors were z-scored along the time dimension, and ridge regression (*α* = 1) was applied at each time window. Statistical assessment followed the same permutation framework as described above. For each participant, a null distribution of regression coefficients was generated by 1000 repeats of circularly shifting the LDA prediction time windows.

### Principal component analysis

Principal component analysis (PCA) reduced source data from the number of source vertices to the first 10 principal components (PCs). Top 3 dimensions are used for visualisation and further analyses. PCA-transformed time series were first smoothed using a one-dimensional Gaussian filter (*σ* = 20 ms) trial-wise and then averaged within each sequence condition.

#### Kolmogorov–Smirnov distance

To quantify the separation between preparation and execution activity along each principal component (PC), we computed the Kolmogorov–Smirnov (KS) distance between their respective one-dimensional distributions. The KS distance is a standard non-parametric measure of distributional dissimilarity, ranging from 0 (identical empirical distributions) to 1 (maximal separation of their empirical cumulative distributions).

For each PC, we performed 1,000 bootstrap iterations on the sequence-averaged PCA trajectories shown in the visualisations. In each iteration, 1,000 time points were randomly sampled from the preparation and execution epochs across the four sequences, and the KS distance between the corresponding PC-value distributions was computed. KS distances were then averaged across iterations.

The null distribution was constructed by repeating the same procedure with randomly permuted phase labels. KS distances were averaged across bootstrap repetitions for each participant, and group-level differences between observed and null values were assessed using paired Wilcoxon signed-rank tests with BH-FDR correction over 3 PCs × 6 areas.

#### Euclidean distance

Separation between preparatory and execution states in the 3D low-dimensional PCA space was quantified as the mean Euclidean distance between preparatory points and the execution centroid. In each bootstrap iteration, let 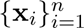 be preparation coordinates and 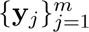 be execution coordinates, the distance is computed as:

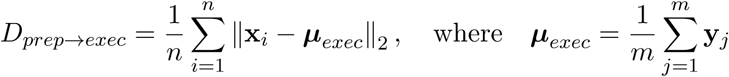

Distances were estimated using the bootstrap procedure (1,000 iterations; 1,000 sampled time points per phase). Null values were generated by permuting phase labels. Group-level observed-versus-null comparisons used paired Wilcoxon signed-rank tests with Bonferroni across 6 regions.

#### Alignment index

The dynamics were first centered by subtracting the cross-condition mean: averaging the condition means across the preparatory and execution phases. To assess the alignment between preparatory and execution subspaces, separate PCAs were performed on the dynamics of each phase (Elsayed et al., 2016). Alignment was then measured by projecting the execution subspace onto the preparatory subspace and measuring the extent of overlap:

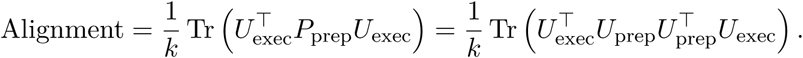

where *U*_prep_ and *U*_exec_ are the PCA basis matrices (with orthonormal columns) for the preparatory and execution subspaces, respectively, and 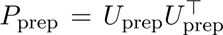 is the projection operator onto the preparatory subspace. A value of 1 indicates perfect alignment, while 0 indicates orthogonality. Statistical significance was assessed by comparing the alignment index against 1 using a one-sample Wilcoxon signed-rank test.

#### Gaussian mixture model and rotation angle

Apart from the subspace alignment index, we also calculated the geometric angle between the two phases. To quantify the angle between preparatory and execution dynamics, separate one-component three-dimensional Gaussian mixture models were fitted to the sequence-averaged PCA embeddings from the preparation and execution phases. These models provided a compact description of the geometric structure of activity within each phase. For each fitted distribution, the principal direction was defined as the eigenvector associated with the largest covariance eigenvalue, corresponding to the direction of greatest variance. The angle between preparatory and execution dynamics was then calculated as the angle between their respective principal directions. Eigenvectors were sign-aligned by enforcing a positive first component to ensure directional consistency across phases and participants.

Angles between states were computed using the dot-product–based acute angle measure:

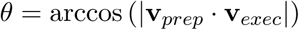

yielding angles constrained to 0^◦^ *−* 90^◦^. For each region, one-sample Wilcoxon signed-rank tests were used to assess whether preparation–execution angles differed from no rotation (0^◦^) and orthogonality (90^◦^) (Bonferroni correction over 6 areas).

### Covariance matrices and t-distributed stochastic neighbour embedding

To visualise temporal changes in covariance structure, we performed a sliding-window covariance analysis on data downsampled to 200 Hz, using a 200 ms window and a 20 ms step. For each participant, covariance matrices were computed across trials and averaged within each sequence and z-scored over time within each sequence. The resulting covariance matrices were then vectorised and embedded into a low-dimensional space using t-distributed stochastic neighbour embedding (t-SNE) (perplexity = 5, ndim = 3). Visualisation was performed in 2D space.

Two summary measures were derived from the embedded covariance trajectories. The mean consecutive Euclidean distance was computed by taking the Euclidean distance between successive sequence-averaged embeddings in the 3-dimensional space. The Euclidean distance between preparation and execution phases was quantified using a bootstrap (1000 repeats): in each iteration, an equal number of samples per phase (N = 1000) was drawn, and the Euclidean distance between the execution-state points and the centroid of the preparation-state points was calculated.

#### Pattern distance between phases

To quantify similarity in covariance structure between preparatory and execution periods, trial-wise covariance matrices were first labelled and grouped according to phase identity. Similarity was estimated using a bootstrap procedure applied separately for each participant. In each of 1,000 bootstrap iterations, 10,000 samples were drawn from each phase and averaged. Covariance similarity was quantified using pattern distance:

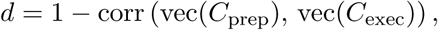

where *C*_prep_ and *C*_exec_ denote the phase-averaged covariance matrices; each is normalised to correlations. vec(*·*) refers to vectorisation of the upper triangular matrix. A null distribution was generated by repeating the full procedure after randomly permuting phase labels, thereby removing the systematic preparatory–execution structure while preserving covariance magnitude and sampling statistics.

### Demixed PCA

Demixed principal component analysis (dPCA) was applied to separate sources of variance attributable to time, sequence identity, and their interaction within the 200 Hz source-reconstructed time series (Kobak et al., 2016). For each participant and each brain region, dPCA was performed on correct trials, using the minimum number of correct trials available across sequences to ensure balanced input. The dPCA input had dimensionality n_sequence_ *×* n_repeats_ *×* n_time_, and the decomposition yielded three sets of principal components corresponding to the time, sequence, and time *×* sequence marginalizations. For each marginalisation, explained variance was quantified and the variance of the top three components was summed.

### Sliding-window LDA for sequence decoding

Sliding-window linear discriminant analysis (LDA) was performed using a 200-ms window with a 10-ms step on the 200 Hz source-level time series. For each participant, 40 correct trials per sequence were randomly sampled without replacement (except P02 used 15 trials per sequence), and data were split into training and test sets using a 70/30 stratified split. Within each time window, the LDA classifier was trained on the corresponding training-set segment and evaluated on the matching held-out test segment. Decoding accuracy was computed as the proportion of correctly classified test-set samples and then averaged across all time points within the window.

This sampling and decoding procedure was repeated 500 times per participant to estimate robust average decoding performance. Final accuracy time courses were obtained by averaging across repetitions. A Gaussian smoothing kernel with *σ* = 40 ms was applied to the curve before both permutation testing and visualisation. All analyses were performed independently for each brain region.

Statistical significance was assessed using a cluster-based permutation test relative to the permuted baseline (cluster-forming threshold = .01, cluster-level significance = .05). For each participant, a null distribution was generated by repeating the decoding procedure with randomly permuted sequence labels (500 permutations). Observed decoding accuracy was then compared against this null distribution to determine whether performance exceeded chance.

The same Gaussian smoothing (*σ* = 40 ms) was applied to both the LDA curves and the frequency-band amplitude curves before performing the regression analysis.

### Sliding-window LDA for factor decoding

To assess discriminability between the two sequence-design factors, we applied the same sliding-window LDA using a 200-ms windows advanced in 10-ms steps. The four learned sequences were collapsed into two binary contrasts reflecting the 2 *×* 2 factorial structure: sequences (1,2) and (3,4) shared the same inter-press interval structure, whereas sequences (1,3) and (2,4) shared the same finger order.

To ensure that training and test sets did not share the same fractal cue, we used two-fold cross-validation across fractal cues (Kornysheva & Diedrichsen, 2014; Yewbrey & Kornysheva, 2024). For the timing contrast, one fold was trained on sequences *{*1, 3*}* and tested on *{*2, 4*}*, with the assignments reversed in the second fold. For the finger-order contrast, one fold was trained on *{*1, 2*}* and tested on *{*3, 4*}*, again with the assignments reversed in the second fold. Within each fold, all subsampled trials from the training sequences were used for fitting, and all trials from the test sequences were used for evaluation. When *n* = 40, each fold contained 40 trials per class for both training and testing. Fold-wise accuracies were averaged across folds and 500 independent resampling repeats.

A label-shuffled null distribution was generated using the same procedure, with 500 repeats per contrast. Within each cross-fractal fold, class labels were randomly permuted among training trials while test labels remained unchanged. This preserved the cross-fractal split structure while disrupting the trial–label relationship. Before plotting and statistical testing, each participant’s accuracy time course was smoothed using a Gaussian kernel with *σ* = 40 ms.

## Results

To dissociate preparatory from execution-related neural patterns that support memory-guided sequential movements across motor and hippocampal circuits, we analysed the MEG dataset reported by Kornysheva et al. (2019).

### Behavioural performance

The experiment used a delayed finger-sequence production task in which participants generated a five-element sequence from memory associated with an abstract visual Sequence Cue (Figure 1A). After a variable preparation interval of 1.8, 2.0, or 2.2 s, a Go Cue prompted sequence execution on a response keyboard. Preparation was defined as the interval from Sequence Cue onset to the Go Cue, and execution as the interval from the first to the last finger press (Figure 1B). Behavioural performance on memory-guided trials improved across training (mean ± SD; day 1: 69.0 ± 22.3%; day 2: 85.7 ± 14.9%; day 1 vs. day 2: p < .001) and remained high during the MEG session (89.3 ± 15.5%; day 2 vs. day 3 MEG: p = .036) (Figure S1A). Behavioural performance did not differ across MEG recording blocks (Friedman test: *χ*^2^(9) = 7.70*, p* = .565; Figure S1B). During the MEG session, mean finger order accuracy was 97.29 *±* 3.43%. Of these, 8.01 *±* 12.30% had the correct order but incorrect timing relative to the target timing structure (Kornysheva et al., 2019), resulting in a mean overall sequence accuracy of 89.3 *±* 15.5% across participants (Figure S1C-E).

**Figure 1:**
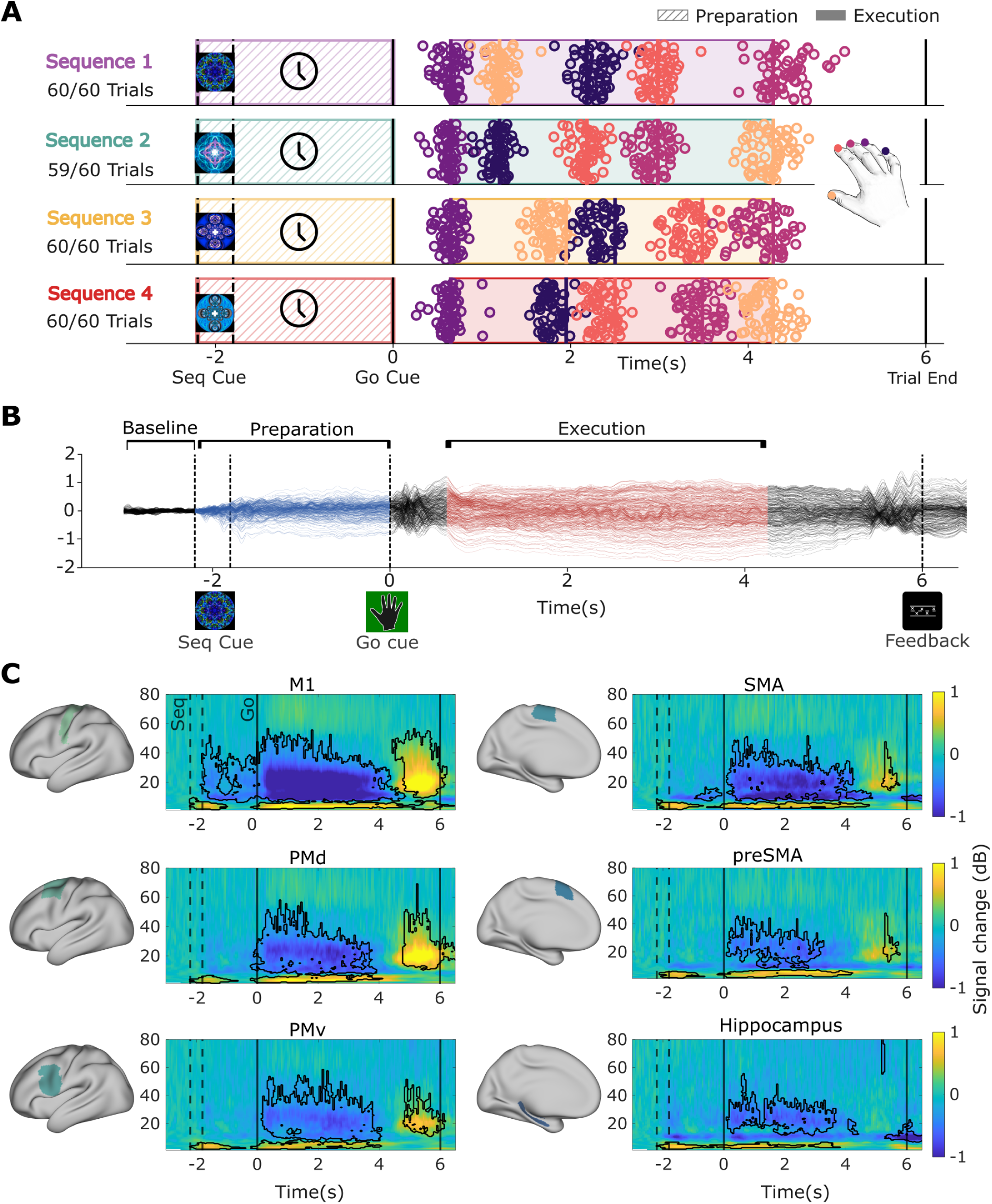
Delayed finger sequence task and time-frequency representations. (A) Task design and behavioural data from a representative participant. The sequences formed a factorial combination of two temporal interval patterns and two finger orderings. During the recording session, each sequence was performed 60 times. Only trials classified as correct in timing and order were shown and included in subsequent analyses. The behavioural data are aligned to the Go Cue; two black dashed vertical lines mark the variable delay period (1.8, 2.0, or 2.2 s) between the Sequence Cue and the Go Cue. Raster dots represent individual finger presses on each trial, colour-coded by finger identity (see fingertip colours in the hand inset). The coloured vertical bars indicate the planned timing of each finger press in the memorised sequence. (B) Overview of the task time course and MEG channel activity averaged across trials and participants. Blue shading indicates the preparation phase between the Sequence Cue and the Go Cue, whereas red shading indicates the execution phase between the first and last finger presses. (C) Time-frequency representations (TFRs) showing source-localised oscillatory dynamics from six regions of interest: M1, PMd, SMA, pre-SMA, PMv and hippocampus. Each panel is paired with the anatomical mask used for source reconstruction. Individual TFRs were baseline-corrected using decibel (dB) normalisation relative to each participant’s own baseline period, then group-averaged and baseline demeaned. Black contours indicate significant time–frequency clusters relative to baseline.

### Oscillatory markers of motor excitation, inhibition and serial recall converge in M1

To localise neural activity from regions of interest across the motor hierarchy reported in our recent fMRI studies involving a similar paradigm (Yewbrey & Kornysheva, 2024; Yewbrey et al., 2023), we applied linearly constrained minimum variance (LCMV) beamforming to source-reconstruct activity from the contralateral primary motor cortex (M1), dorsal premotor cortex (PMd), ventral premotor cortex (PMv), supplementary motor area (SMA), pre-supplementary motor area (pre-SMA), and the hippocampus (Figure 1C).

Consistent with prior work, we detected signatures of movement-related neural excitation and inhibition (Barone & Rossiter, 2021), including beta- and alpha (mu)-band event-related desynchronisation(ERD) during sequence preparation and execution, followed by a post-movement beta rebound (PMBR; Figure 1C). Preparatory ERD was significant in M1, where it emerged approximately 1.85 seconds before the Go Cue and persisted through the end of execution. This pattern was followed by smaller ERD clusters in premotor regions, including PMd, PMv, SMA, and pre-SMA (Figure 1C). Cluster-level p-values and the associated time and frequency ranges are provided in Table 1.

**Table 1:**
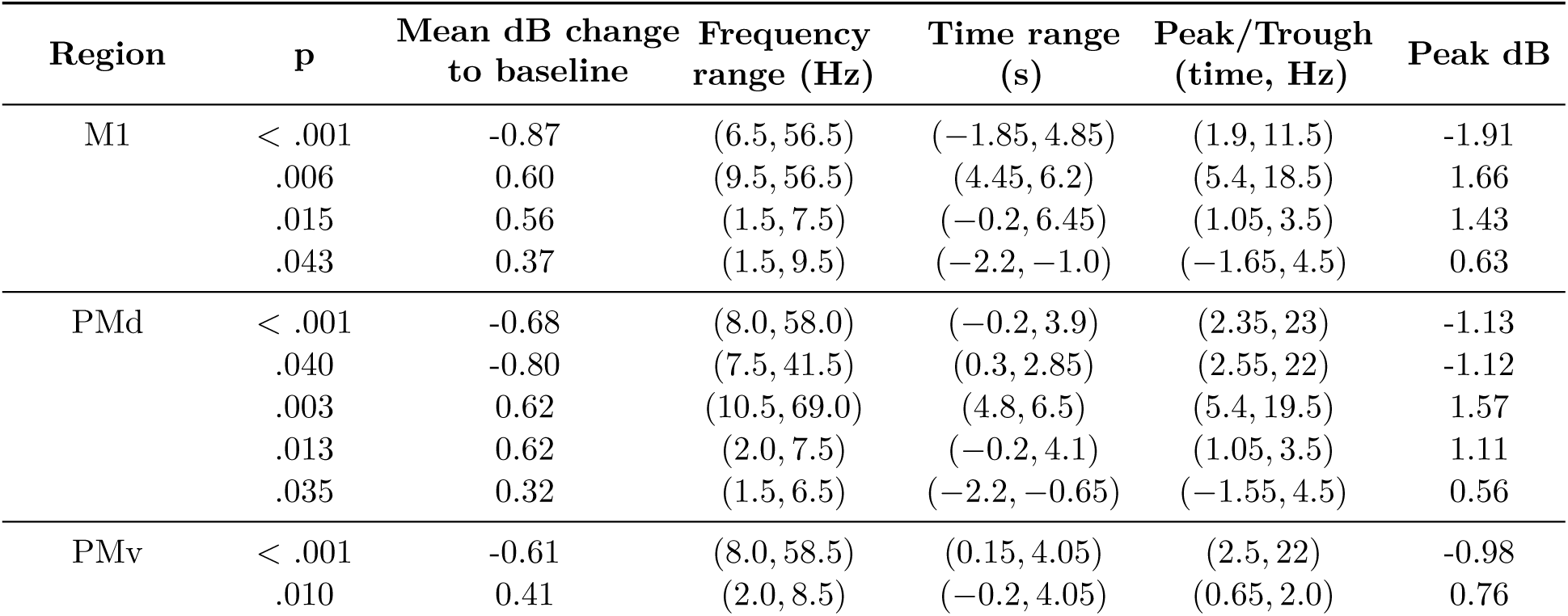
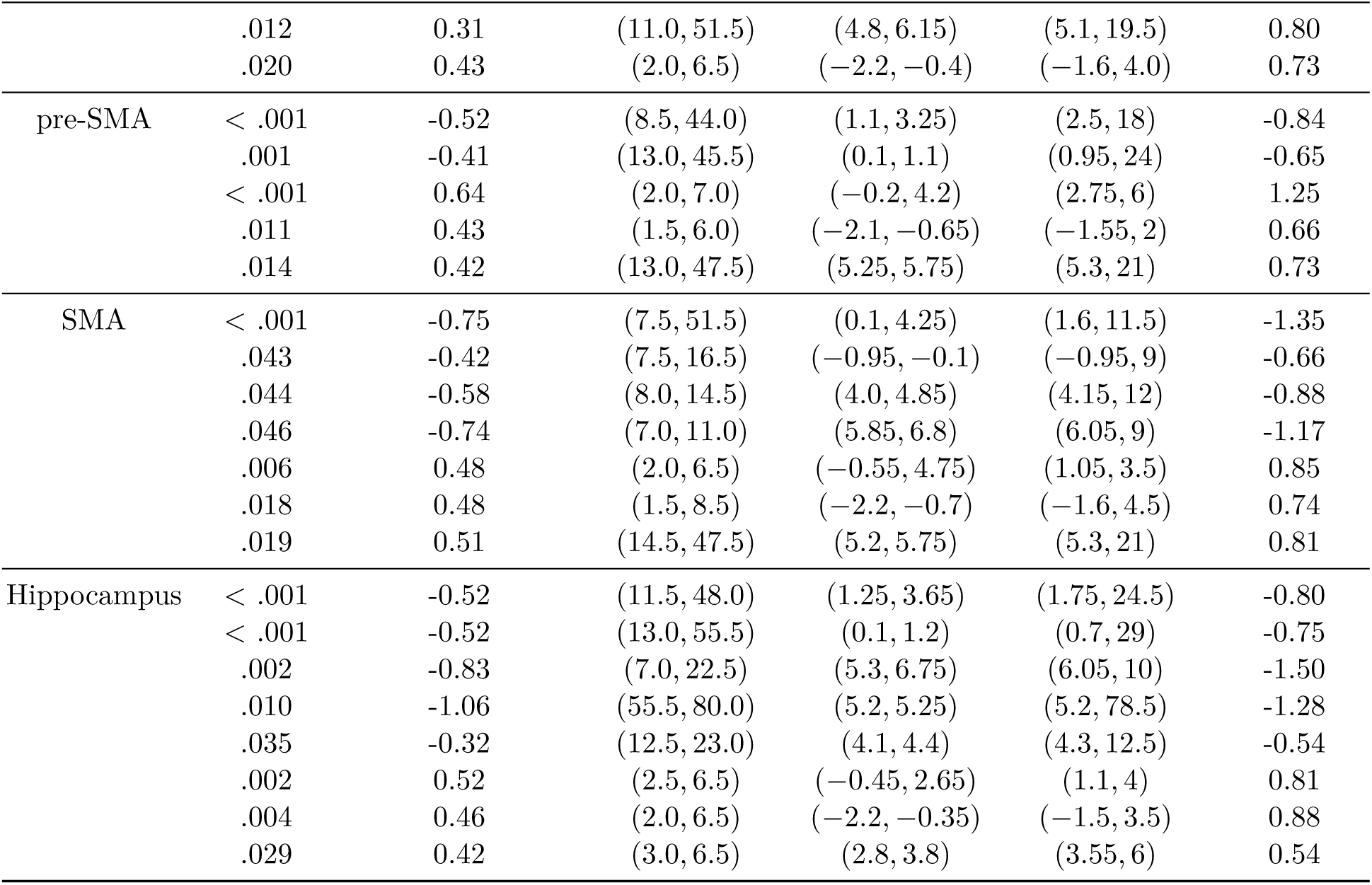
Significant time–frequency clusters by region.

| Region | p | Mean dB change to baseline | Frequency range (Hz) | Time range (s) | Peak/Trough (time, Hz) | Peak dB |
| --- | --- | --- | --- | --- | --- | --- |
| M1 | < .001 | -0.87 | (6.5, 56.5) | (-1.85, 4.85) | (1.9, 11.5) | -1.91 |
|  | .006 | 0.60 | (9.5, 56.5) | (4.45, 6.2) | (5.4, 18.5) | 1.66 |
|  | .015 | 0.56 | (1.5, 7.5) | (-0.2, 6.45) | (1.05, 3.5) | 1.43 |
|  | .043 | 0.37 | (1.5, 9.5) | (-2.2, -1.0) | (-1.65, 4.5) | 0.63 |
| PMd | < .001 | -0.68 | (8.0, 58.0) | (-0.2, 3.9) | (2.35, 23) | -1.13 |
|  | .040 | -0.80 | (7.5, 41.5) | (0.3, 2.85) | (2.55, 22) | -1.12 |
|  | .003 | 0.62 | (10.5, 69.0) | (4.8, 6.5) | (5.4, 19.5) | 1.57 |
|  | .013 | 0.62 | (2.0, 7.5) | (-0.2, 4.1) | (1.05, 3.5) | 1.11 |
|  | .035 | 0.32 | (1.5, 6.5) | (-2.2, -0.65) | (-1.55, 4.5) | 0.56 |
| PMv | < .001 | -0.61 | (8.0, 58.5) | (0.15, 4.05) | (2.5, 22) | -0.98 |
|  | .010 | 0.41 | (2.0, 8.5) | (-0.2, 4.05) | (0.65, 2.0) | 0.76 |

| Region | p | Mean dB change<br>to baseline | Frequency<br>range (Hz) | Time range<br>(s) | Peak/Trough<br>(time, Hz) | Peak dB |
| --- | --- | --- | --- | --- | --- | --- |
|  | .012 | 0.31 | (11.0, 51.5) | (4.8, 6.15) | (5.1, 19.5) | 0.80 |
|  | .020 | 0.43 | (2.0, 6.5) | (-2.2, -0.4) | (-1.6, 4.0) | 0.73 |
| pre-SMA | < .001 | -0.52 | (8.5, 44.0) | (1.1, 3.25) | (2.5, 18) | -0.84 |
|  | .001 | -0.41 | (13.0, 45.5) | (0.1, 1.1) | (0.95, 24) | -0.65 |
|  | < .001 | 0.64 | (2.0, 7.0) | (-0.2, 4.2) | (2.75, 6) | 1.25 |
|  | .011 | 0.43 | (1.5, 6.0) | (-2.1, -0.65) | (-1.55, 2) | 0.66 |
|  | .014 | 0.42 | (13.0, 47.5) | (5.25, 5.75) | (5.3, 21) | 0.73 |
| SMA | < .001 | -0.75 | (7.5, 51.5) | (0.1, 4.25) | (1.6, 11.5) | -1.35 |
|  | .043 | -0.42 | (7.5, 16.5) | (-0.95, -0.1) | (-0.95, 9) | -0.66 |
|  | .044 | -0.58 | (8.0, 14.5) | (4.0, 4.85) | (4.15, 12) | -0.88 |
|  | .046 | -0.74 | (7.0, 11.0) | (5.85, 6.8) | (6.05, 9) | -1.17 |
|  | .006 | 0.48 | (2.0, 6.5) | (-0.55, 4.75) | (1.05, 3.5) | 0.85 |
|  | .018 | 0.48 | (1.5, 8.5) | (-2.2, -0.7) | (-1.6, 4.5) | 0.74 |
|  | .019 | 0.51 | (14.5, 47.5) | (5.2, 5.75) | (5.3, 21) | 0.81 |
| Hippocampus | < .001 | -0.52 | (11.5, 48.0) | (1.25, 3.65) | (1.75, 24.5) | -0.80 |
|  | < .001 | -0.52 | (13.0, 55.5) | (0.1, 1.2) | (0.7, 29) | -0.75 |
|  | .002 | -0.83 | (7.0, 22.5) | (5.3, 6.75) | (6.05, 10) | -1.50 |
|  | .010 | -1.06 | (55.5, 80.0) | (5.2, 5.25) | (5.2, 78.5) | -1.28 |
|  | .035 | -0.32 | (12.5, 23.0) | (4.1, 4.4) | (4.3, 12.5) | -0.54 |
|  | .002 | 0.52 | (2.5, 6.5) | (-0.45, 2.65) | (1.1, 4) | 0.81 |
|  | .004 | 0.46 | (2.0, 6.5) | (-2.2, -0.35) | (-1.5, 3.5) | 0.88 |
|  | .029 | 0.42 | (3.0, 6.5) | (2.8, 3.8) | (3.55, 6) | 0.54 |

PMBR was most pronounced in contralateral M1 and PMd, with progressively weaker modulation in SMA, pre-SMA, and PMv. Several beta-band ERD clusters were also observed in the hippocampus during execution, but no PMBR was detected. Instead, the hippocampus showed a high-gamma ERD cluster after movement termination.

Increases in theta- and delta-band power were observed across all regions during both preparation and execution, in line with their role in memory and serial recall and their expected distribution across both hippocampal and cortical sources (Aktürk et al., 2022; Herweg et al., 2020). These low-frequency clusters were detected in M1, PMd, PMv, SMA, pre-SMA, and the hippocampus.

These time-frequency results confirm the predicted oscillatory markers in motor, premotor, and hippocampal regions during a memory-guided motor sequence task. Further, they indicate that contralateral M1, the principal cortical output area, is pre-activated during sequence preparation and shows all task-related band modulations observed in upstream areas.

### Timing of the transition from preparation to execution patterns

To determine the timing of the neural transition from sequence preparation to execution, we trained and tested an eight-class linear discriminant analysis (LDA) classifier on trial-wise MEG activity patterns separately for each of the six regions. The eight classes represented the four sequences during preparation and execution (Figure 2A).

**Figure 2:**
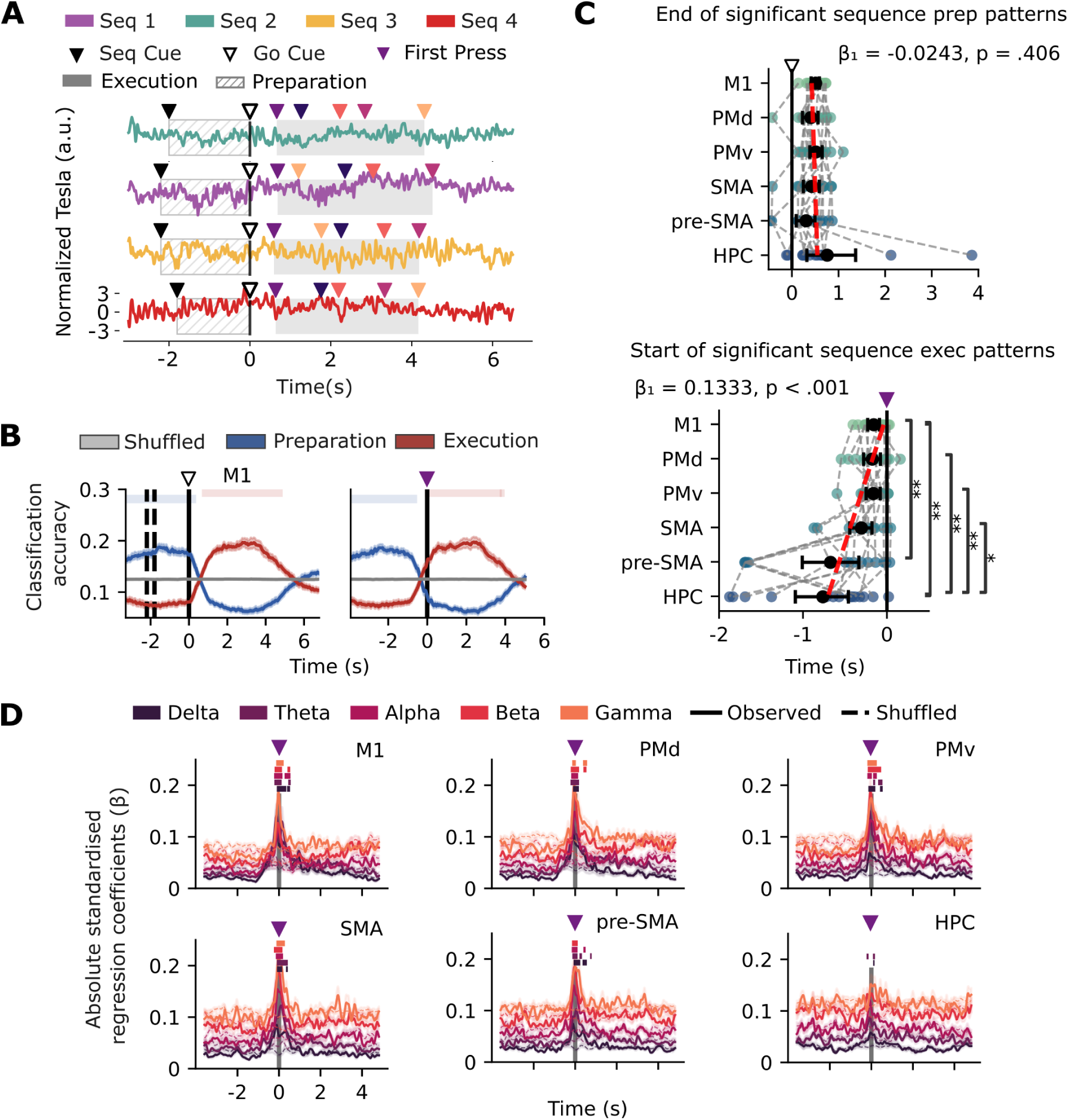
LDA quantifies temporal transitions between motor preparation and execution. (A) Schematic of the eight-class Linear Discriminant Analysis (LDA) training. The classifier discriminated between combinations of sequence identity and task phase (e.g., Seq1–Prep and Seq2–Exec) using 200-Hz source-reconstructed dynamics from individual trials. The schematic shows one trial from each of the four sequences, with trial-specific behavioural markers defining the preparation and execution phases. Equal numbers of samples were drawn from each phase. To address class imbalance, 40 trials were sampled per sequence (except for P02), and results were averaged over 1,000 bootstrap repetitions. (B) LDA classification performance across participants (n = 14) in M1, aligned to the Go Cue (left) or the first button press (right). Curves show the accuracy assigned to the correct sequence–phase class (e.g., Seq1-Prep / Seq1-Exec for sequence-1 trials), averaged across participants; shaded regions denote standard error. Semi-transparent bars at the top of each plot mark significant intervals relative to the shuffled baseline (cluster-forming alpha < .01, final threshold < .05). Preparatory and execution predictions exhibit temporally non-overlapping patterns. (C) Offset of significant preparation patterns, aligned to the Go Cue (top), and onset of significant execution patterns, aligned to the first press (bottom) across motor areas and the hippocampus. Connecting lines represent data from the same participant. Statistical comparisons were performed using Wilcoxon signed-rank tests with BH–FDR correction across 15 comparisons. Red dashed lines show the fitted linear mixed-effects models computed from participant transition timings. Corresponding model outputs are shown in the panels. (D) Rectified standardised regression coefficients relating the execution-classification time course to canonical frequency-band activity, anchored to movement onset. Colours denote frequency bands. Shading at the top of each plot marks significant intervals relative to the permuted-regression baseline (circular-shift null, averaged over 1,000 permutations). Both the execution-probability time course and the TFR-derived band-power signals were z-scored across the whole trial prior to regression. Regressions were computed using a sliding-window (200 ms window, 10 ms step).

If preparation and execution were not separable, the classifier would fail to discriminate phase above chance. If sequence tuning generalised across phases, classifier evidence would transfer between corresponding preparation and execution sequence labels. Alternatively, if preparation and execution relied on phase-specific representations, these should produce the strongest classifier evidence and accuracy for labels matching the sampled phase.

The resulting sequence probability time courses showed distinct and temporally non-overlapping sequence patterns (Figure 2B; Figure S2A-B): execution-related sequence classification accuracy remained below chance during the preparatory period, including shortly after the Go Cue, and rose significantly above chance only shortly before execution. Preparatory tuning showed the opposite pattern, with a brief gap between the phases where neither phase reached significance. This stark result confirmed that the preparation and execution phases are associated with distinct neural sequence tuning patterns that do not generalise across phases. This pattern was consistent across all regions, with varying decoding strength from M1 to hippocampus (Figure S2C).

To resolve the timing of the offset of the preparatory sequence patterns and the onset of the execution patterns above chance, classifier accuracy was aligned to two behavioural anchors: the Go Cue and the first movement (button press). Significance testing against randomly permuted class labels showed that neural transitions consistently occurred after the Go Cue and before movement initiation. The offset of sequence preparation patterns was quantified for each region: M1: 0.51 *±* 0.17 s; PMd: 0.41 *±* 0.32 s; PMv: 0.51 *±* 0.27 s; SMA: 0.42 *±* 0.34 s; pre-SMA: 0.30 *±* 0.40 s; hippocampus: 0.78 *±* 1.04 s (Figure 2C, upper panel). Similarly, the onset of the sequence execution patterns differed across regions: M1: *−*0.16 *±* 0.14 s; PMd: *−*0.18 *±* 0.20 s; PMv: *−*0.16 *±* 0.16 s; SMA: *−*0.30 *±* 0.27 s; pre-SMA: *−*0.68 *±* 0.69 s; hippocampus: *−*0.76 *±* 0.65 s (Figure 2C, lower panel). Post-hoc comparisons revealed significant differences between M1 and pre-SMA (*p* = .0097), as well as between the hippocampus and M1 (*p* = .0097), PMd (*p* = .0097), PMv (*p* = .0097), and SMA (*p* = .032).

To investigate whether the timing of neural pattern transitions followed the motor hierarchy, we fitted linear mixed-effects models. The hierarchical order was defined by the proximity of regions to the periphery, i.e., in line with the reported anatomical density of direct cortico-spinal projections to the lower cervical segments of the spinal cord (C7-T1) and cortico-cortical connectivity to M1 (Dum & Strick, 2005), from furthest to closest: hippocampus, pre-SMA, SMA, PMv, PMd and M1. Whilst the offset of preparation patterns showed no evidence of a hierarchical shift (*β*_0_ = 0.571*, β*_1_ = *−*0.024*, p* = .406), the onset of execution revealed a significant positive slope, with transitions occurring progressively later along the motor hierarchy and M1 transitioning last to execution patterns (*β*_0_ = *−*0.838*, β*_1_ = +0.133*, p* < .001).

Regressing the execution-prediction curve onto spectral power across canonical frequency bands revealed that significant clusters were concentrated around the preparation–execution transition (Figure 2D, Table 2). M1 showed significant regression coefficients across the delta, theta, alpha, beta and gamma bands. Similar broadband regression coefficients were observed in PMd, PMv, SMA and pre-SMA, whereas hippocampal coefficients were restricted to the delta and theta bands.

**Table 2:**
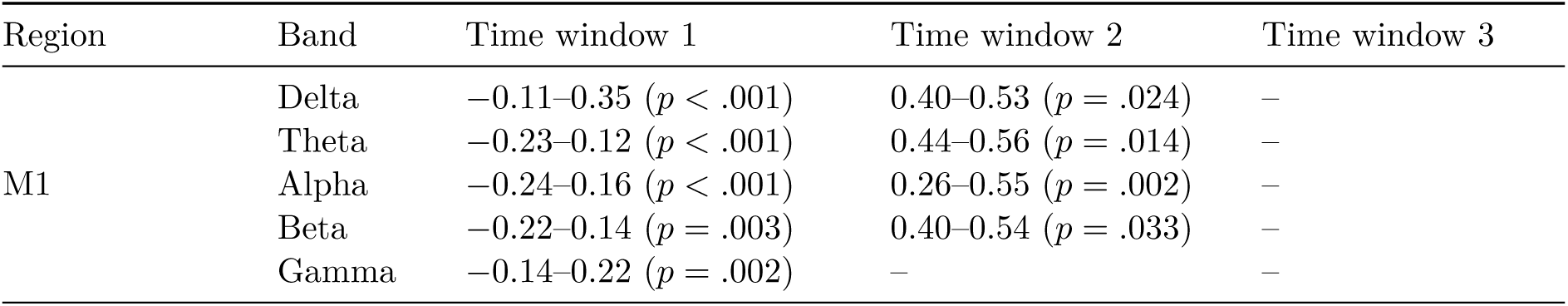
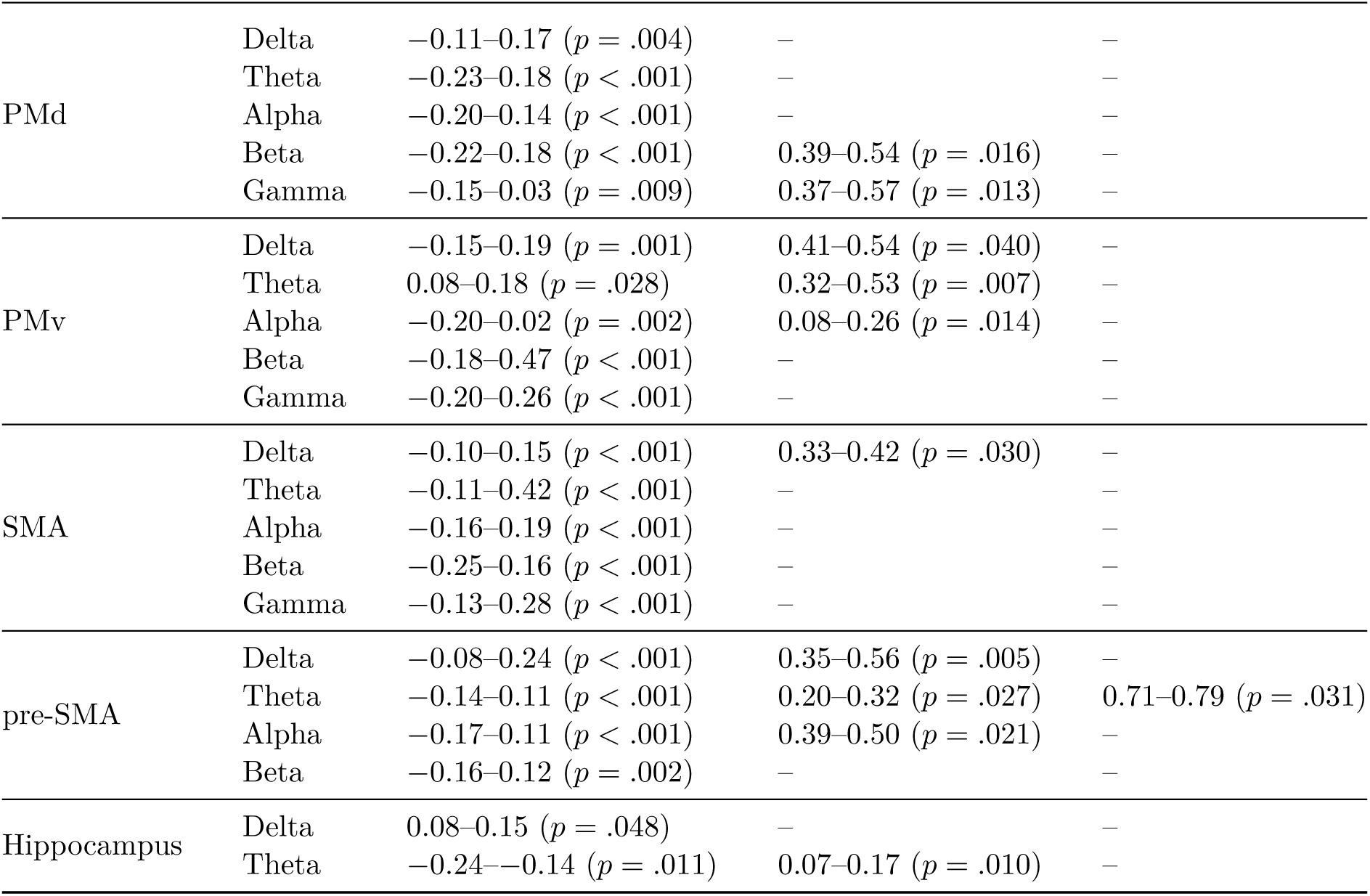
Significant clusters from regressions between execution-state prediction curves and spectral power across canonical frequency bands. Cluster windows are reported relative to movement onset.

### Principal component analysis reveals distinct preparation and execution manifolds

LDA revealed a temporal contrast between the two phases. To investigate whether the population geometry was orthogonal between preparation and execution, as reported in invasive animal studies (Churchland & Shenoy, 2024; Churchland et al., 2012), we applied principal component analysis (PCA) to embed high-dimensional source MEG dynamics into three-dimensional manifolds. The first three principal components accounted for most of the variance across all regions (M1: 90.5 *±* 3.3%; PMd: 94.9 *±* 1.7%; PMv: 93.3 *±* 2.2%; SMA: 97.5 *±* 1.1%; pre-SMA: 97.7 *±* 0.7%; hippocampus: 99.2 *±* 0.3%), thereby enabling quantification of geometric properties in the low-dimensional space.

When projected into this low-dimensional space, trajectories from the preparation and execution phases occupied distinct manifolds, showing differential distributions between phases along the principal component axes (Figure 3A). This phase separation was confirmed by statistically significant Kolmogorov–Smirnov (KS) distances between the preparation and execution distributions along each PC axis, evaluated against a permutation-based null distribution and present in all regions of interest (Figure 3B; all *p* < .001; Wilcoxon signed-rank tests, FDR corrected across 6 areas × 3 PCs).

**Figure 3:**
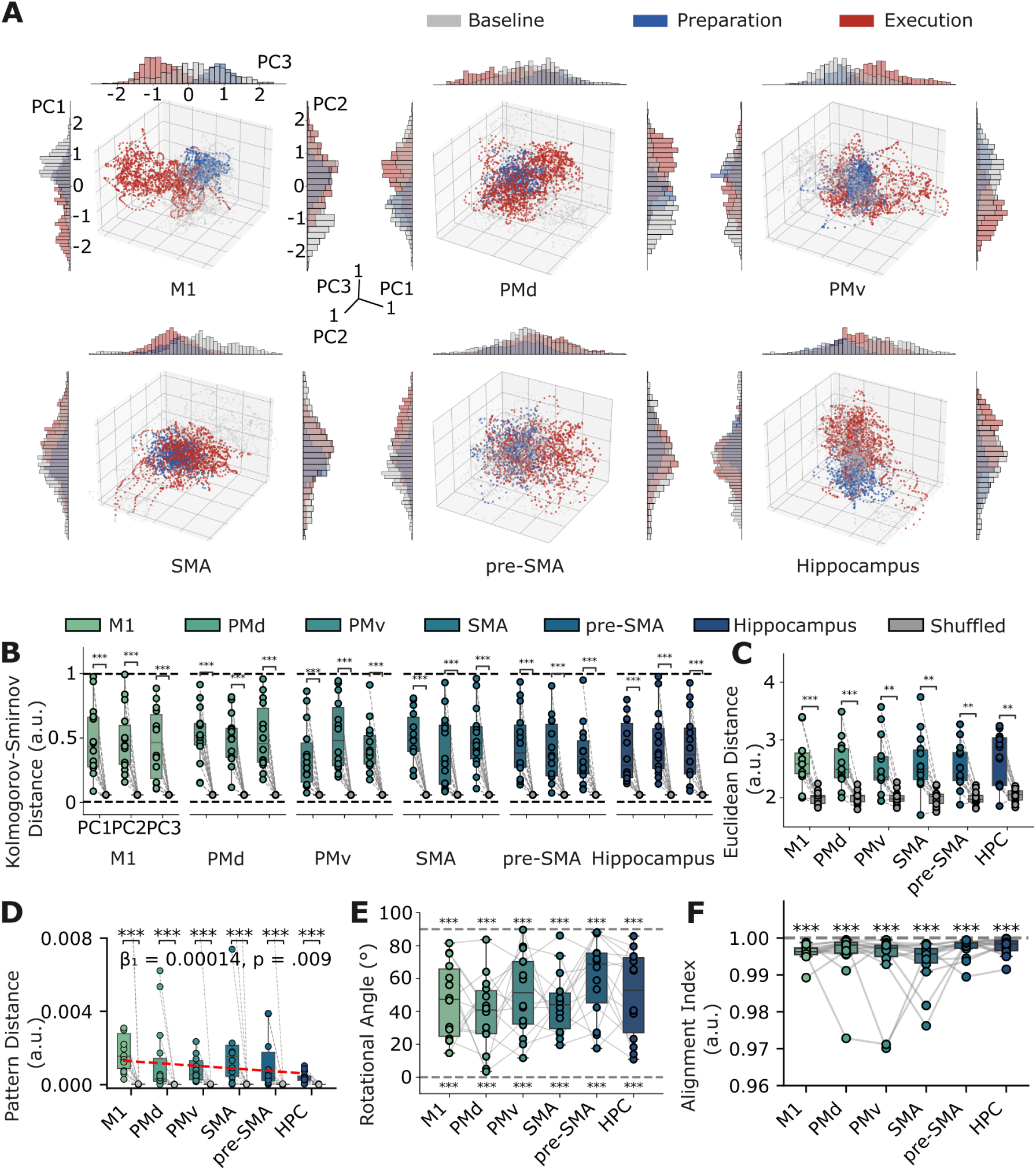
Preparation and execution occupy distinct subspaces in a shared low-dimensional manifold. (A) Source-localised MEG data projected onto the first three principal components for the representative participant. All axes were z-scored. Trajectories show sequence-averaged dynamics, colour-coded by phase (preparation, execution). Marginal histograms indicate the distribution of data points along each principal component axis. (B) Kolmogorov–Smirnov (KS) distances between preparatory and execution dynamics along individual principal components. The KS distance (range: 0–1) quantifies the maximum difference between the empirical distributions of two samples. For each participant, KS distances were estimated by bootstrapping 1000 data points over 1,000 iterations. Significance was assessed against a null distribution generated by randomly permuting phase labels. (C) Euclidean distance between preparatory and execution trajectories in the 3D PCA space. For each participant, the mean distance from execution points to the preparatory centroid was compared to a null distribution obtained by randomly permuting phase labels (1000 samples per group, 1,000 iterations). (D) Pattern distance between phase-average covariance matrices. Time-windowed covariance matrices were sampled within each phase, averaged to reduce within-phase variability, and compared between preparation and execution (1000 bootstraps). Distances were significantly greater than chance across all regions (Wilcoxon signed-rank tests against permutation; Bonferroni-corrected across 6 regions; p < .001). (E) Angle between the directions of maximum variance of the preparatory and execution Gaussian mixture models (GMMs). This angle reflects the difference in orientation between the two dominant axes of variation. The resulting angles were neither parallel (0^◦^) nor orthogonal (90^◦^) to each other (Wilcoxon signed-rank tests, Bonferroni-corrected across 6 regions; *p* < .001 for all comparisons). (F) Alignment index quantifying the similarity between neural manifolds formed by preparatory and execution dynamics. Values approaching 1 indicate stronger alignment, suggesting that both phases occupy similar subspaces. Wilcoxon signed-rank test against full alignment (1) yielded *p* < .001 for all areas (Bonferroni correction over 6 areas).

Furthermore, the Euclidean distance between the phase clusters in the low-dimensional space, which indicates the degree of geometric separation between the two phases, was significantly greater than chance in all regions (Figure 3C; corrected across 6 regions; M1: 2.31 *±* 0.43; PMd: 2.30 *±* 0.42; PMv: 2.27 *±* 0.45; SMA: 2.30 *±* 0.48; pre-SMA: 2.27 *±* 0.39; hippocampus: 2.36 *±* 0.46; *p ≤ .*0015 for all regions). Although M1 and the hippocampus showed the largest mean separations, these differences were not statistically significant across regions (Friedman test: *χ*^2^(5) = 2.12, *p* = .832), indicating that the degree of phase separation was comparable across areas in PCA space.

To directly test whether the preparatory and execution periods exhibit distinct or similar covariance patterns, we computed covariance matrices in a sliding-window fashion and quantified the similarity distance between the preparatory and execution matrices. The correlation metric is scale-invariant; therefore, the distance between phases indicates that execution is not simply a weaker or scaled version of preparation. In all regions, the pattern distance between preparatory and execution covariance matrices was significantly higher than the shuffled equivalent (Figure 3D; tests against permutation; corrected across 6 regions; *p* < .001 for all regions). The Friedman test indicated a modest effect of brain areas, *χ*^2^(5) = 11.14, *p* = .049, but no pairwise comparisons survived correction across 15 area pairs. However, a linear mixed-effects model revealed a significant increase in pattern distance along the anatomically ordered hippocampus-to-M1 gradient (*β*_0_ = 0.000480*, β*_1_ = 0.000138*, p* = .009).

Previous research highlighted the geometric orthogonality between the preparatory and execution dynamics (Kaufman et al., 2014), whereby distinct principal components selectively capture activity from the two phases. In a three-dimensional projection, this organisation would produce approximately orthogonal orientations of the phase-specific variance. Conversely, substantial sharing of principal components between the two phases would be expected to produce more closely aligned orientations. To examine this in our data, we first fitted a Gaussian mixture model (GMM) to each state within the three-dimensional PCA space and compared the angles between their respective principal axes (Figure 3E). The resulting angles were neither parallel (0^◦^) nor orthogonal (90^◦^) to each other (M1: 46.7 *±* 23.1^◦^; PMd: 38.4 *±* 22.8^◦^; PMv: 51.2 *±* 24.9^◦^; SMA: 45.1 *±* 19.2^◦^; pre-SMA: 60.6 *±* 24.0^◦^; hippocampus: 50.0 *±* 26.3^◦^; Wilcoxon signed-rank tests, Bonferroni-corrected across 6 regions; *p* < .001 for all comparisons).

Complementary to the rotation analysis, which evaluates the orthogonality of the two phases within a shared low-dimensional embedding, we performed a subspace-alignment analysis (Elsayed et al., 2016) to quantify the overlap between subspaces derived independently from preparatory- and execution-related dynamics. Orthogonal subspaces yield minimal overlap and, consequently, a low alignment index, indicating that execution-related variance cannot be represented within the preparatory subspace. The alignment index was high, indicating that the preparatory and execution dynamics occupy highly similar low-dimensional manifolds (Figure 3E; M1: 0.997 *±* 0.0019; PMd: 0.994 *±* 0.011; PMv: 0.993 *±* 0.0098; SMA: 0.993 *±* 0.0066; pre-SMA: 0.998 *±* 0.0018; Hippocampus: 0.999 *±* 0.0010). However, when tested against perfect alignment (index = 1), all areas showed alignment indices significantly below 1 (Wilcoxon signed-rank tests, Bonferroni-corrected across 6 regions; all *p* < .001). A Friedman test indicated a regional effect (*χ*^2^(5) = 13.10; *p* = .022), but no pair-wise comparison survived FDR-BH correction.

### MEG covariance pattern separation is driven by phase, not sequence identity

Whereas the PCA analyses characterised the geometric relationship between preparatory and execution-related activity, covariance analysis examined how the underlying correlation structure changed over time. To capture this temporal evolution, we computed covariance matrices in sliding windows, quantifying the time-resolved co-variation and spatial coordination among local neural populations across the preparation-to-execution transition. These high–dimensional matrices were then embedded into a low-dimensional space by t-distributed stochastic neighbour embedding (t-SNE), providing a sensitive probe of phase- and sequence-specific neural coding dynamics across brain regions during motor behaviour (Figure 4A).

**Figure 4:**
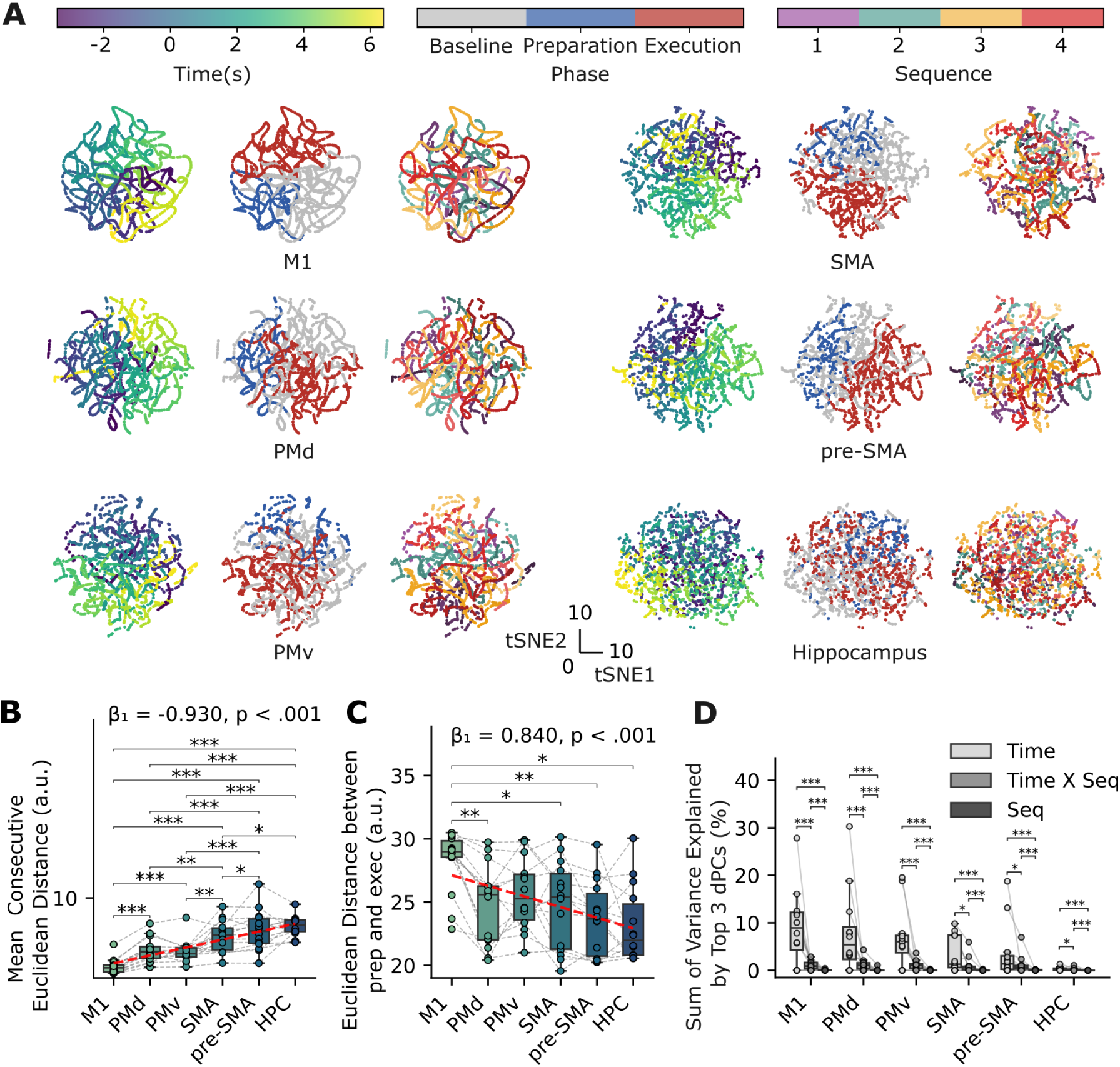
Time-resolved covariance structure reveals phase-specific clustering. (A) t-distributed stochastic neighbour embedding (t-SNE) of time-resolved covariance matrices from six regions of interest (M1, PMd, PMv, SMA, pre-SMA, and hippocampus) in an example participant. Covariance matrices were computed using 200 Hz binned data with a 200 ms sliding-window and a 20 ms stepsize and embedded into a 2D t-SNE space. Each subpanel consists of three small plots showing continuous time label (left), phase label (middle) and sequence label (right). (B) Mean consecutive Euclidean distance between adjacent time points in the t-SNE space, each dot is one participant. Smaller distances indicate smoother, more temporally continuous trajectories, whereas larger distances reflect discontinuous or irregular transitions. Horizontal brackets denote significant pairwise differences (Wilcoxon signed-rank test, BH–FDR corrected over 15 comparisons). The red line indicates the fitted linear mixed-effects model. (C) Euclidean distance between preparatory and execution clusters within the t-SNE embedding, quantifying phase separation in each region. M1 exhibits high separation, indicating a sharper transition between preparatory and execution states. Individual participant values are shown with paired lines. Horizontal brackets denote significant pairwise differences (Wilcoxon signed-rank, BH–FDR corrected). The red line indicates the fitted linear mixed-effects model. (D) Variance explained by the top three demixed PCA (dPCA) components for each region, decomposed into contributions from time, sequence, and time *×* sequence factors. Each dot represents one participant; boxplots show group distributions. Asterisks mark significant differences between component types within region (Wilcoxon signed-rank tests, BH–FDR corrected).

Critically, this analysis highlighted heterogeneity in how different brain regions traversed this phase transition: the primary motor cortex (M1) showed the greatest temporal continuity, reflected by the smallest consecutive distances (Figure 4B; *mean ± SD*; M1: 1.97 *±* 0.90; PMd: 3.88 *±* 1.45; PMv: 3.73 *±* 1.36; SMA: 5.54 *±* 1.84; pre-SMA: 6.21 *±* 2.37; Hippocampus: 6.72 *±* 1.18). In contrast, the hippocampus displayed a more scattered and discontinuous trajectory, indicating weaker internal organisation or sequential changes in local coordination relative to the task phases. A Friedman test confirmed robust regional differences in trajectory continuity (*χ*^2^(5) = 60.204, *p* < .001). Post-hoc Wilcoxon signed-rank tests with BH-FDR across 15 area pairs revealed significant differences (the following pairs had *p* < .001: M1 vs. PMd, M1 vs. PMv, M1 vs. SMA, M1 vs. pre-SMA, M1 vs. hippocampus, PMd vs. SMA, PMd vs. pre-SMA, PMd vs. hippocampus; PMv vs. pre-SMA, PMv vs. hippocampus; other significant pairs: PMv vs. SMA: *p* = .003; SMA vs. hippocampus: *p* = .021). This regional trend was confirmed by a linear mixed-effects model, which revealed a significant negative slope across the hippocampus-to-M1 hierarchy (*β*_0_ = 7.929, *β*_1_ = *−*0.930, *p* < .001), indicating greater temporal continuity toward M1.

Next, we quantified the distance between low-dimensional phase clusters, which provides a measure of preparatory-to-execution progression (Figure 4C; *mean ± SD*; M1: 28.25 *±* 2.43; PMd: 24.70 *±* 3.21; PMv: 25.62 *±* 2.80; SMA: 24.72 *±* 3.49; pre-SMA: 23.73 *±* 2.96; Hippocampus: 23.14 *±* 3.01). M1 showed significantly greater phase separation than PMd, SMA, pre-SMA and hippocampus (Friedman test: *χ*^2^(5) = 16.490, *p* = .006; post-hoc Wilcoxon signed-rank tests, BH-FDR corrected over 15 comparisons: M1 vs. PMd: *p* = .005; M1 vs. SMA: *p* = .011; M1 vs. pre-SMA: *p* = .005; M1 vs. Hippocampus: *p* = .011), suggesting a structured, continuous remapping of spatial coding underlying motor control. A linear mixed-effects model confirmed a significant positive trend in phase separation along the hierarchical ordering (*β*_0_ = 22.089, *β*_1_ = 0.840, *p* < .001), with greater separation toward M1. This suggests that phase transitions are not merely governed by scaling a fixed pattern, but rely on gradual, structural shifts in spatial neural coding.

To quantify this statistically, we sought to dissociate the variance arising from time-dependent changes and sequence identity by applying demixed principal component analysis (dPCA; Figure 4D). This method extends PCA by constraining components to separately capture variance attributable to time, sequence identity, or their interaction. The majority of variance was aligned with the temporal transition, with the hippocampus exhibiting the least variance and M1 showing the highest variance among motor regions (M1: 9.13 *±* 7.97%; PMd: 7.87 *±* 8.99%; PMv: 6.85 *±* 6.42%; SMA: 3.25 *±* 3.78%; pre-SMA: 3.69 *±* 5.97%; Hippocampus: 0.34 *±* 0.38%). In contrast, components specifically attributed to sequence identity accounted for only a small proportion of the total variance and showed weaker separability across sequences (Figure 4D; time x sequence: M1: 1.04 *±* 0.90%; PMd: 1.16 *±* 1.21%; PMv: 1.12 *±* 1.03%; SMA: 1.08 *±* 1.71%; pre-SMA: 1.19 *±* 1.93%; Hippocampus: 0.19 *±* 0.30%; sequence: M1: 0.07 *±* 0.11%; PMd: 0.10 *±* 0.31%; PMv: 0.05 *±* 0.07%; SMA: 0.04 *±* 0.07%; pre-SMA: 0.06 *±* 0.07%; Hippocampus: 0.01 *±* 0.01%). Within each region, Wilcoxon tests across the three marginalisations (BH-FDR, 18 comparisons) were significant for all contrasts. Fifteen had *p* < .001; the largest adjusted p-values were for time versus time × sequence in pre-SMA (*p* = .024), SMA (*p* = .036), and hippocampus (*p* = .042).

To examine whether the variance explained by the sequence-related components differed across regions, we performed Friedman tests across ROIs. Significant regional differences were observed for both the sequence × time component (Friedman test: *χ*^2^(5) = 24.76, *p* < .001) and the sequence component (*χ*^2^(5) = 21.90, *p* < .001) (Figure 4D). Post-hoc Wilcoxon tests with BH-FDR (15 comparisons per marginal) showed higher variance in motor regions than in the hippocampus for both marginals (sequence × time: M1 (*p* = .007), PMd (*p* = .002), PMv (*p* = .002), SMA (*p* = .002), and pre-SMA (*p* = .002); sequence: M1 (*p* = .004), PMd (*p* = .004), PMv (*p* = .013), and pre-SMA (*p* = .013).

### Decoding sequence identity

Finally, we set out to test whether sequence-specific patterns, whilst distinct across phases, could be used to decode sequence identity. We applied sliding-window LDA, training and testing within the same peri-movement phase, to probe when sequence identity could be recovered during preparation and production (Figure 5A). Using short, 200-ms windows aligned to equivalent time points within each phase reduced the influence of phase-transition-related variance and isolated sequence-specific information.

**Figure 5:**
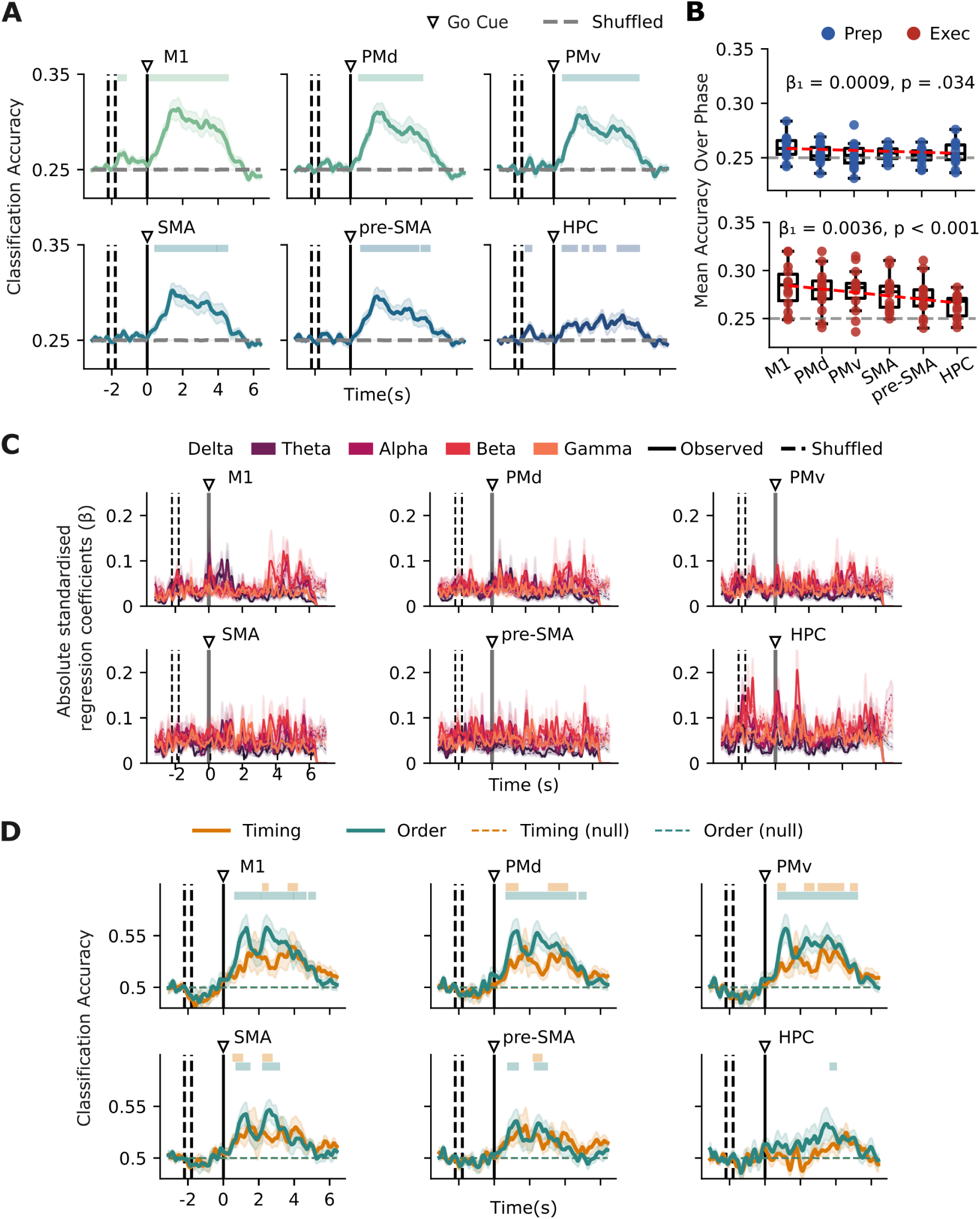
Sliding-window decoding of sequence identity. (A) Sliding-window LDA decoding of sequence identity. A linear discriminant classifier was trained and tested within each time window (200 ms window, 10 ms step) to decode the four finger sequences. Curves were smoothed with a Gaussian kernel with *σ* = 40 ms for both permutation and visualisation. Solid lines indicate participant averaged classification accuracy based on real labels; grey lines represent chance-level performance(n = 14); shading indicates SD. Top bars mark time intervals where decoding exceeded shuffled baseline (cluster-forming alpha = .01, final cluster alpha = .05). All motor areas showed above-chance decoding during execution, with region-specific timing differences. The hippocampus exhibited lower decoding accuracy. (B) Sliding-window accuracy averaged over the preparation window and execution window. The red line indicates the fitted linear mixed-effects model. (C) Regression coefficients relating frequency band power to decoder output. Coloured lines represent the standardised regression coefficient of delta (1–4 Hz), theta (4–8 Hz), alpha (8–13 Hz), beta (13–30 Hz), and gamma (30–80 Hz) bands. Coloured shading at the top indicates time points with significant regression relative to the shuffled regression null distribution (cluster-forming alpha = .01, final cluster alpha = .05, Bonferroni-corrected across frequency bands). (D) Sliding-window LDA decoding of sequence factors. A sliding-window LDA classifier was trained and tested within each time window (200 ms window, 10 ms step) to decode either temporal or ordinal design.

Decoding accuracy significantly exceeded the shuffled baseline, primarily during execution in all motor regions (Figure 5B). M1 reliably recovered sequence identity during sequence preparation (-1.64 to -1.2 s, p = .025) together with a brief window in the hippocampus (-1.58 to -1.26 s, p = .044). We then averaged decoding accuracy across the preparation and execution periods for each region and tested for hierarchical trends using linear mixed-effects models. Significant hierarchical effects were observed during both preparation (*β*_0_ = 0.251, *β*_1_ = 0.0009, *p* = .034) and execution (*β*_0_ = 0.263, *β*_1_ = 0.0036, *p* < .001).

To evaluate the spectral contributions to decoding, trial-wise LDA outputs were regressed onto z-scored power in canonical frequency bands (Figure 5C). The resulting regression coefficients were temporally unstable, and most did not reach significance relative to the shuffled regression baseline, suggesting that no individual frequency band reliably drives sequence decoding.

To determine whether the neural data contained information about the ordinal or temporal features of the sequences, we performed analogous sliding-window decoding analyses for each factor separately. Specifically, we tested whether the classifier could distinguish sequences that differed in finger order or in timing pattern (Figure 5D). Decoding was generally stronger during execution than during preparation, with finger-order information producing higher accuracy than temporal-pattern information. In the hippocampus, finger-order information was decodable, whereas no significant decoding window was observed for the temporal factor.

### Primary visual cortex as a non-motor control region

To evaluate the functional and regional specificity of our findings, we conducted a set of post-hoc control analyses in the primary visual cortex (V1). V1 served as an optimal non-motor control region given its lack of direct cortico-spinal output and exhibited the expected stimulus-evoked profile: alpha / beta desynchronization across preparation and execution, together with cue-aligned gamma-power increases (Figure S4A; Table S1). This profile provided an independent benchmark for the source-reconstructed time–frequency analysis.

First, analyses of phase-pattern separation (Figure S2A-B) and neural population geometry (Figure S6A) revealed that the separation between preparatory and execution manifolds was not restricted to motor regions and the hippocampus but also extended to V1. This finding suggests that preparation-to-execution state shifts represent a widespread cortical principle rather than an isolated motor property — a dynamic that may have been overlooked in animal models due to the historical sampling bias toward M1 and PMd (Kaufman et al., 2014; Zimnik & Churchland, 2021).

Second, although V1 exhibited distinct preparatory and execution manifolds, its transition to an execution-like population state occurred late and was strictly time-locked to movement onset, whereas the hippocampus transitioned well beforehand, followed sequentially by pre-SMA (Figure S5A). This temporal dissociation confirms that the early state shift in the hippocampus reflects a specific preparatory mechanism rather than a generalised timing artefact of non-motor cortical processing.

Third, V1 exhibited a significantly lower alignment index between preparatory and execution subspaces than M1 (p = .018), PMd (p = .045), pre-SMA (p = .021), and the hippocampus (p = .018; Figure S6F). However, the alignment index did not indicate complete separation between the subspaces, and the inter-phase rotation angle did not differ significantly across regions (Figure S6E). Thus, V1 did not exhibit fully orthogonal preparatory and execution manifolds. The partial overlap observed in V1 highlights that strict orthogonality may be difficult to resolve in non-invasive, source-reconstructed signals, which can reflect mixtures of underlying neural populations. Alignment and rotation metrics therefore provide a graded characterisation of representational similarity and separation, rather than relying on complete orthogonality as an all-or-none criterion.

Fourth, the geometry of V1 neural representations more closely resembled that of M1 than that of higher-order regions, including the hippocampus, pre-SMA, and SMA. V1 showed small consecutive-window Euclidean distances in the tSNE embedding, comparable to M1 but significantly smaller than those observed in the other regions (Figure S7B). Preparation–execution distances in the t-SNE embedding were also comparable between V1 and M1 but differed significantly from those observed in the higher-order regions (Figure S7C).

Finally, multivariate pattern decoding revealed that V1 supported robust above-chance sequence classification during the preparation period, exceeding the decoding accuracy observed in M1 and all other ROIs (all p < .001; Figure S8A-B). This high preparatory classification is consistent with sensory encoding of the fractal sequence cues, which remained on screen during preparation and held a unique, participant-specific mapping to each sequence. Significant regressions between sequence decoding and spectral power across canonical frequency bands around Sequence Cue onset further support the interpretation that V1 was strongly engaged by visual input (Figure S8C; Table S3). Decoding of the factors was absent during preparation (Figure S8D), providing a negative control and arguing against memory-driven decoding in V1. Unexpectedly, V1 also sustained high decoding accuracy during sequence execution, which was driven more strongly by spatial finger-press order than by sequence timing. Because participants’ peripheral vision was unobstructed while they maintained central fixation, this late execution decoding likely reflects peripheral visual feedback during manual button pressing.

In summary, while V1 participated dynamically in visual cue processing and exhibited broad preparation-to-execution manifold shifts, its late transition timing and state-space geometry sharply distinguish it from high-level hubs in the sequence control hierarchy, such as the hippocampus and pre-SMA.

## Discussion

### State change from preparation to execution

Invasive neural recordings in humans and non-human primates reveal a consistent computational principle during the peri-movement phase: cortical motor areas maintain orthogonal population dynamics to functionally separate motor preparation from motor execution, preventing premature motor output (Churchland et al., 2012; Dekleva et al., 2024; Elsayed et al., 2016; Kaufman et al., 2014; Russo et al., 2020). Our MEG results extend this notion beyond primarily motor regions in the context of preparation and execution of skilled motor sequences in humans. We show that the neural activity underlying motor sequence preparation and execution from memory undergoes a pronounced global state shift prior to movement onset across motor, premotor, medial prefrontal and hippocampal regions previously shown to be involved in memory-guided motor sequence control (Yewbrey & Kornysheva, 2024; Yewbrey et al., 2023). This fundamental geometric property of neural dynamics ensures the decoupling of planning and acting, whether the motor output is a discrete single action or part of a complex, memory-retrieved behavioural sequence. Importantly, these findings demonstrate that the distinction between preparation and execution manifolds is not restricted to motor and premotor areas, but extends to distributed regions across the brain, including those involved in visual task processing. This result suggests that the state shift between preparation and execution may represent a broader, system-wide dynamic across cortical networks – a critical insight overlooked by previous electrophysiological studies that focused on manifold dynamics during single movements and sequences of movements using recordings restricted to small subsets of motor regions, such as M1 and premotor cortices, evaluated in isolation (Churchland & Shenoy, 2024; Kaufman et al., 2014; Zimnik & Churchland, 2021).

### Hierarchical transition to execution patterns

Execution-related sequence patterns did not unfold simultaneously, in contrast to the synchronous termination of preparatory patterns, reset by the visual Go Cue. Instead, their onset varied systematically across regions before the movement, consistent with a hierarchical organisation and the regions’ proximity to the periphery (M1 and motor units in the spinal cord) (Dum & Strick, 2005). In the hippocampus, pre-SMA, and SMA, the shift toward execution patterns appeared *∼*600 ms or more and *∼*300 ms before the first button press, respectively, whereas in M1, an area closer to the periphery due to dense cortico-spinal outputs, the shift emerged only *∼*160 ms before finger press onset. Although this temporal ordering aligns with a top–down progression from higher- to lower-level motor-related areas close to the periphery, the *∼*500 ms delay between the state changes in pre-SMA and M1 cannot be explained by direct cortico-cortical interactions (Alexander et al., 1986; Civardi et al., 2001; Ferraina et al., 2002; Hikosaka et al., 2002; Mars et al., 2009). This points toward an iterative involvement of subcortical, likely striato-thalamo-cortical loops in triggering state changes across brain regions (Logiaco et al., 2021).

Despite evidence for hierarchically timed transitions to execution states, we observed substantial inter-individual variability in the timing of these transitions, particularly in premotor areas and the hippocampus, with some participants showing state transitions to execution-driven patterns even before the Go Cue onset. This could reflect differences in task strategy, such as different offsets of abstract sequential cognitive planning versus onsets of lower-level kinaesthetic motor imagery when motor sequences are retrieved from memory and prepared for execution, albeit in the absence of movement.

Across primary motor and premotor cortices, the oscillatory signatures underlying this state transition were not confined to a single frequency band, such as the beta or alpha modulation observed during motor preparation and production, but were instead broadband. In contrast, the hippocampus exhibited a band-specific profile, relying primarily on theta activity to predict the transition. Motor state transitions from preparation to execution recruit multiple physiological processes, such as the release of inhibition reflected in beta activity, increases in excitability indexed by alpha (mu) desynchronisation over motor areas (Barone & Rossiter, 2021; Van Wijk et al., 2012), and local processing often associated with higher-frequency activity resulting in broadband spectral changes (Fries, 2009). The hippocampus, in contrast, supports sequence retrieval and memory-guided planning, functions tightly coupled to theta oscillations. Theta organises hippocampal firing sequences (phase precession, theta sequences) (Buzsáki & Moser, 2013) and is the dominant rhythm for these computations. Accordingly, hippocampal state transitions are predominantly mediated by theta-dependent phase coding, which is sufficient to represent shifts between mnemonic or planning states (Ólafsdóttir et al., 2018).

### Sequence representations are phase-dependent and dynamic

Whilst the neurophysiological patterns associated with sequence preparation and execution occupied distinct subspaces, suggesting limited overlap in sequence tuning, they were not completely orthogonal and showed significant covariance. This residual overlap suggests that preparing an entire movement sequence inherently recruits components of the execution-related representation. Notably, this stands in contrast to prior findings in single-movement tasks, where preparatory and execution states have been shown to be more cleanly dissociable (Kaufman et al., 2014). One possibility is that motor sequence preparation engages processes akin to motor imagery, whereby aspects of the upcoming movements are internally simulated before execution. Such internal simulation — particularly in the form of kinaesthetic motor imagery (Stinear et al., 2006) — could account for the partial overlap observed between preparatory and execution-related neural states. This overlap may be related to the lack of sequential movement fusion, as the presses, although well trained, were not retrieved as one holistic movement, but as a concatenation of kinematically discrete elements (Spencer & Ivry, 2005; Wright-Wieckowski et al., 2026).

Neural patterns related to the trial phase before and after movement onset accounted for the largest portion of variance in MEG dynamics (dPCA), whereas time and sequence identity had a smaller influence. Notably, whole-sequence patterns did not generalise across the peri-movement phase, indicating that sequence-specific coding changes dynamically between motor sequence preparation and execution. In other words, activity does not occupy a separate “space” for each sequence; rather, it evolves collectively before and after movement onset, yet each sequence retains its own distinctive trajectory within those subspaces. The most pronounced distance between sequences unfolds in the execution phase when movements diverge in their identity and timing after the first press for execution. The trajectories exhibited regional differences in continuity with smoother trajectories in M1 than in upstream motor regions. This echoes population trajectories recorded invasively during sequential motor control in non-human primate M1 versus SMA (Russo et al., 2020).

Although the sequential code was subtle and highly dynamic, MEG patterns still allowed reliable decoding of the motor sequences during execution across all areas and even before movement onset in the hippocampus and M1. While decoding accuracy was modest and far from sufficient for non-invasive BCI applications, it is nevertheless striking that sequential information can be extracted from non-invasive MEG dynamics, given how similar the finger press sequences were: they involved the same digits, pairs sharing the same digit order but differing in temporal structure, the same number of presses and an identical first digit across sequences within each sequence set per participant. Unlike previous single-cell and fMRI studies reporting premotor tuning to full movement sequences during preparation (Ariani et al., 2025; Shima & Tanji, 2000; Tanji & Shima, 1994; Yewbrey et al., 2023), we found no above-chance decoding during preparation in premotor areas, with widespread decoding only during execution. Further, decoding of sequence order and timing could not be detected above chance during preparation, in contrast to previous fMRI studies suggesting the presence of generalisable patterns during preparation (Yewbrey & Kornysheva, 2024; Yewbrey et al., 2023), which suggests that preparatory and execution sequence representations rely on distinct neurophysiological regimes: sequence planning may engage asynchronous, highly temporally jittered spatial ensemble coding (visible to fMRI but invisible to MEG), whereas execution transforms these plans into phase-synchronised, large-scale population oscillations that produce robust MEG signals. This discrepancy might be resolved via cross-modal fusion analyses (Cichy & Oliva, 2020).

Decoding accuracy was not found to be related to power changes across multiple oscillatory frequencies. This suggests that sequence information is not primarily driven by global markers of neuronal excitation and inhibition, such as beta event-related desynchronisation or rebound in motor areas. Instead, decoding performance is likely driven by richer temporal and phase-related neural dynamics beyond simple spectral power changes.

### Limitations and future directions

While both MEG and invasive microelectrode arrays provide millisecond temporal resolution, they capture fundamentally distinct neurophysiological substrates. MEG measures extracellular magnetic fields generated by synchronised postsynaptic currents across large pyramidal populations (Baillet, 2017), whereas invasive arrays resolve single-unit action potentials and local field potentials within micro-circuits. Consequently, MEG signals decay rapidly with depth and are biased toward superficial, fissural cortical architecture (Hillebrand & Barnes, 2002), introducing spatial blurring and potential source leakage between proximate cortical nodes. Despite these biophysical constraints, our findings demonstrate that non-invasive macro-scale population trajectories reliably capture the low-dimensional manifold rotations separating motor preparation from execution. This indicates that key state-space properties established in animal electrophysiology are embedded within human macro-scale signal topologies and accessible without invasive recording.

Evaluating primary visual cortex (V1) as a non-motor control region provides crucial boundaries regarding the functional specificity and regional distribution of state-space dynamics. Historically, preparatory and execution manifolds have been examined almost exclusively within motor and premotor structures (Churchland & Shenoy, 2024; Kaufman et al., 2014; Zimnik & Churchland, 2021). We show that V1, a task-relevant but non-motor region, also reorganises into distinct preparatory and execution subspaces—driven by visual cue processing and peripheral visual feedback during keypresses. Crucially, however, V1’s state transition occurred strictly at movement onset (Figure S5A), sharply contrasting with the prospective, pre-movement transition observed in the hippocampus. Thus, while V1 participated dynamically in visual processing, its manifold structure aligned with lower-level execution dynamics rather than high-level preparatory hubs (hippocampus and pre-SMA). This temporal dissociation between non-motor regions confirms that while state-space reorganisation reflects a distributed cortical response to shifting task phases, MEG can successfully isolate top-down, sequence-specific planning dynamics from low-level sensorimotor processing.

Several methodological considerations qualify these findings. First, our paradigm investigated consolidated, memory-guided sequence control, where motor planning is intrinsically coupled with memory retrieval and cue-sequence association (Kornysheva et al., 2019). While trial-phase jittering and dual-anchoring analyses (locking to the Go Cue and movement onset) ensured that observed state shifts were specifically tied to movement initiation, future studies using non-memory control tasks are needed to further dissociate general motor preparation from memory-guided sequence retrieval. Second, our sample size (*N* = 14), though typical for dense electrophysiological paradigms, represents a limitation that constrains statistical power for detecting subtle effects or higher-order factor interactions.

More broadly, our results highlight the advantage of whole-brain non-invasive electrophysiology. Emerging technologies such as optically pumped magnetometer MEG (OPM-MEG) (Boto et al., 2018), combined with advanced source reconstruction, enable simultaneous mapping of population dynamics across distributed cortical and subcortical networks without the field-of-view restrictions of invasive electrode arrays. Accessing these multi-regional interactions non-invasively will be vital for next-generation brain–computer interfaces (BCIs) capable of interpreting cognitive intent and complex motor sequences prior to movement execution (Andersen et al., 2019; Gallego et al., 2022; Tanji & Shima, 1994).

### Conclusions

In sum, we show that motor sequence preparation and execution involve a global state shift across motor, premotor and hippocampal regions, unfolding hierarchically from areas distant from the periphery to those with dense corticospinal connections. Preparatory and execution-related patterns are largely distinct but partially overlapping, consistent with internal simulation or motor imagery of upcoming sequences. Sequence-specific information is subtle, dynamic, and distributed across multiple frequencies and phases, emerging during preparation only in the hippocampus and M1 (alongside V1), and most strongly across regions during execution but detectable even before movement onset. Critically, non-invasive MEG captures these population-level dynamics, highlighting both fundamental principles of hierarchical motor control and their relevance for brain–computer interfaces, where monitoring multiple regions beyond M1 could enhance decoding of user intent.

## Author Contributions

Ziwei Yin: Formal analysis; Investigation; Methodology; Software; Visualization; Writing—Original draft; Writing—Review and editing. Jian Liu: Conceptualization; Formal analysis; Funding acquisition; Methodology; Software; Supervision; Writing—Review and editing. Katja Kornysheva: Conceptualization; Formal analysis; Funding acquisition; Methodology; Resources; Supervision; Writing—Original draft; Writing—Review and editing.

## Acknowledgments

This work was supported by a UKRI Future Leaders Fellowship awarded to Katja Kornysheva (MR/Y016467/1). This work was partially supported by BlueBEAR and Baskerville, funded by the EPSRC and UKRI through the World Class Labs scheme (EP/T022221/1) and the Digital Research Infrastructure programme (EP/W032244/1) and operated by Advanced Research Computing at the University of Birmingham.

## Data Availability

Researchers may access the study data and materials by email to the lead author.

## Citation Diversity Statement

The authors of this article report its proportions of citations by gender category to be: M/M = 0.58; W/M = 0.24; M/W = 0.1; W/W = 0.08. This is in line with the rate of publications in broad-scope neuroscience journals, but underciting M/W and W/W gender categories relative to the rate published in the Journal of Cognitive Neuroscience (https://psych.wisc.edu/gcbialyzer/).

## Appendix: Supplementary Figures

**Figure S1:**
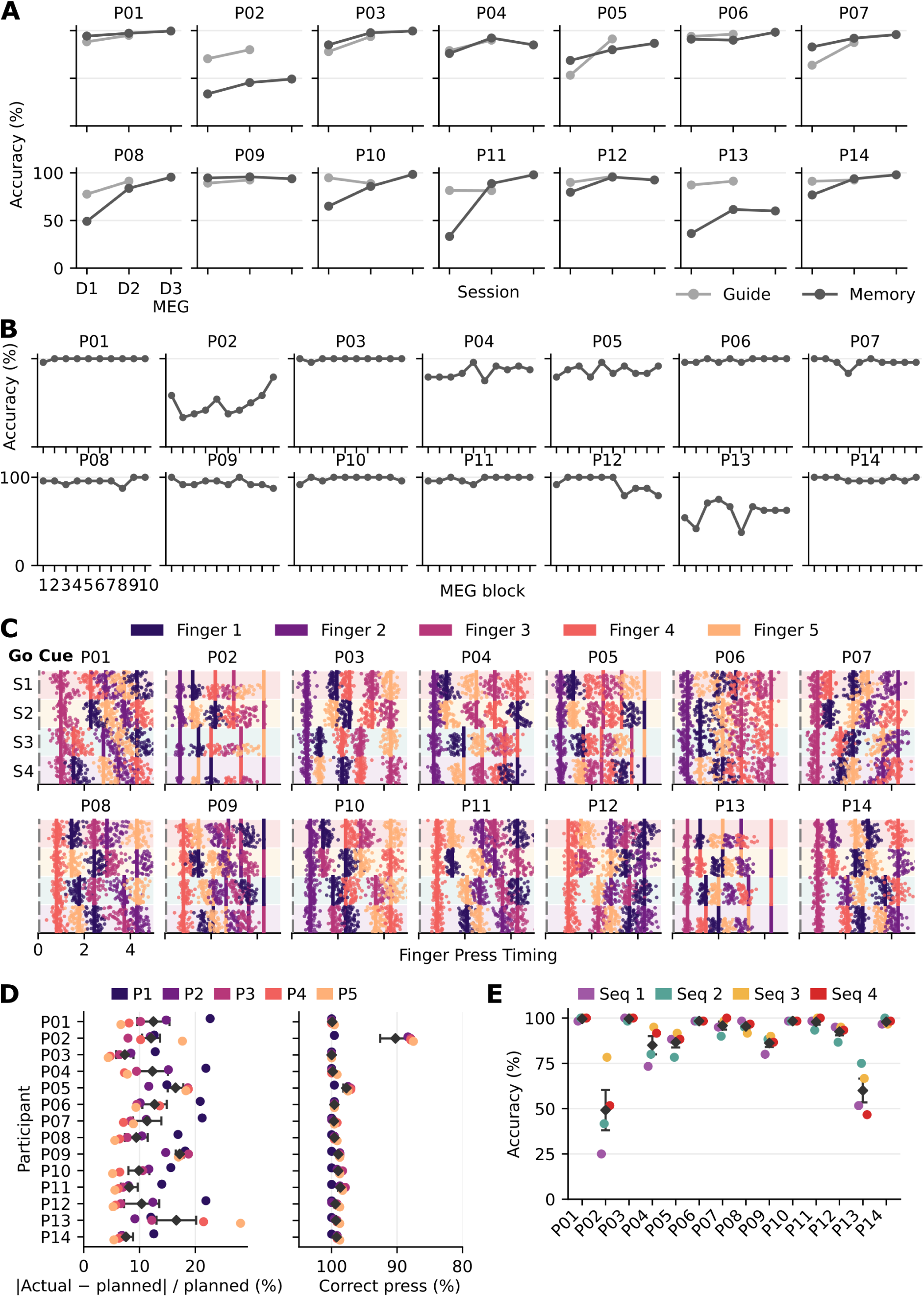
Behavioural data during training and MEG sessions. (A)Participant-level accuracy across training days and the MEG session, shown separately for instructed (Guide) and memory-guided (Memory) trials. (B) Participant-level accuracy across MEG recording blocks. Each block contained 6 trials per sequence, with 10 blocks in total. (C) Raster plots of finger presses across all participants. Only trials correct in both order and timing are shown. Each row represents one correct trial, with each point indicating a finger press and its colour denoting finger identity. Vertical lines indicate the planned timing of each press. (D) Timing and finger-order accuracy for participants. The left panel shows relative timing error for each finger position, calculated as the absolute percentage difference between the observed and planned press times, divided by the planned press time. The right panel shows the percentage of presses with the correct finger order at each sequence position. Points are coloured by press position. (E) Participant-level sequence accuracy shown as a dot plot. Coloured points represent accuracy for Sequences 1–4. Error bars represent uncertainty across trials.

**Figure S2:**
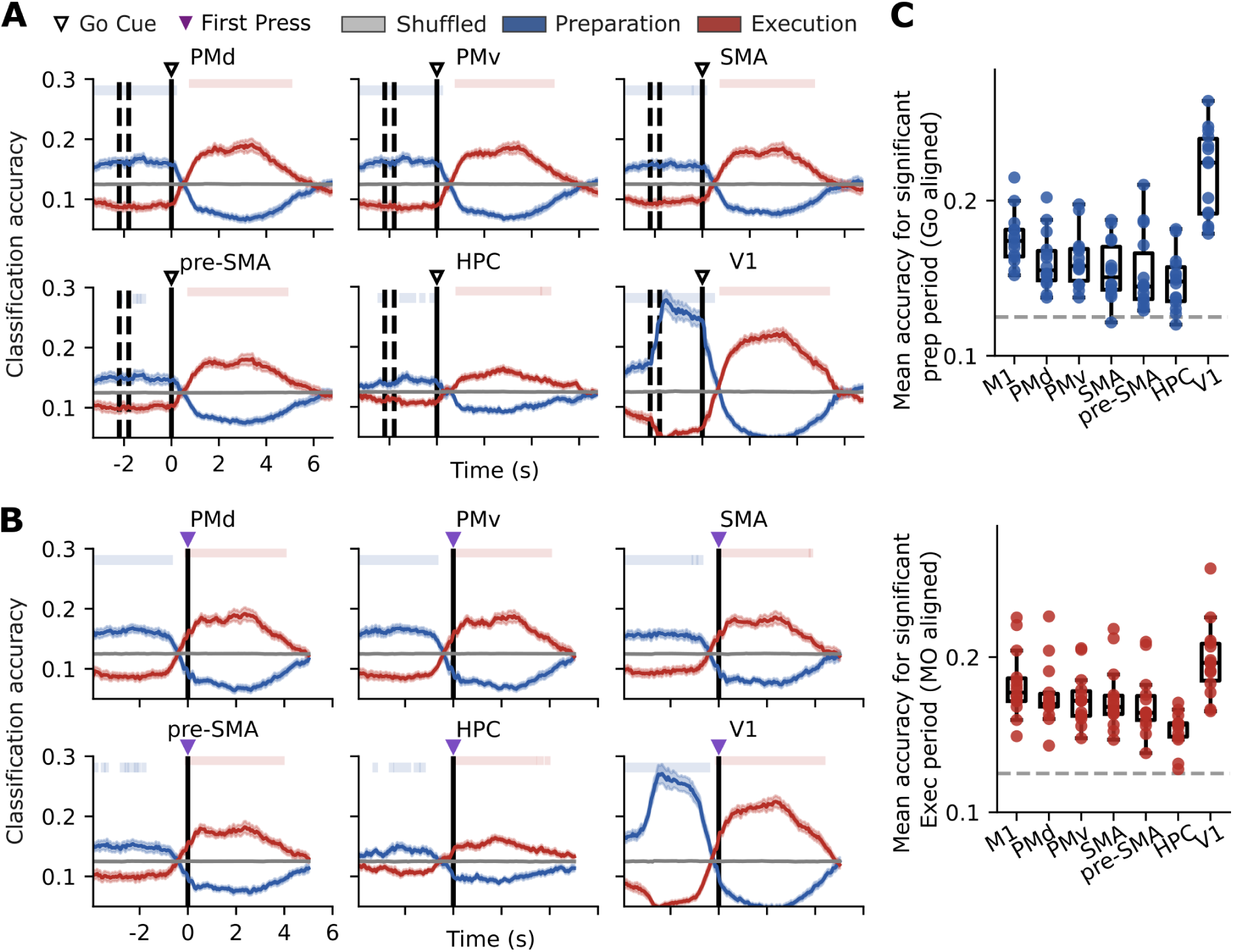
Accuracy and decoding dynamics for the phase LDA. (A) Time-resolved classification accuracy aligned to the Go Cue. Curves show participant-average, with shaded regions indicating the standard error. The open triangle marks the Go Cue. Background shading denotes the preparation and execution periods; the grey line indicates chance level. Top bars in both panels indicate time points at which decoding was significantly above the permutation-derived chance level. (B) The same analysis aligned to the first finger press. The purple triangle marks the first press. (C) Mean classification accuracy within the significant temporal clusters for the preparation period aligned to the Go Cue (top) and the execution period aligned to the first finger press (bottom). Individual points represent participants, boxplots summarise the group distributions. The dashed grey line indicates chance level.

**Figure S3:**
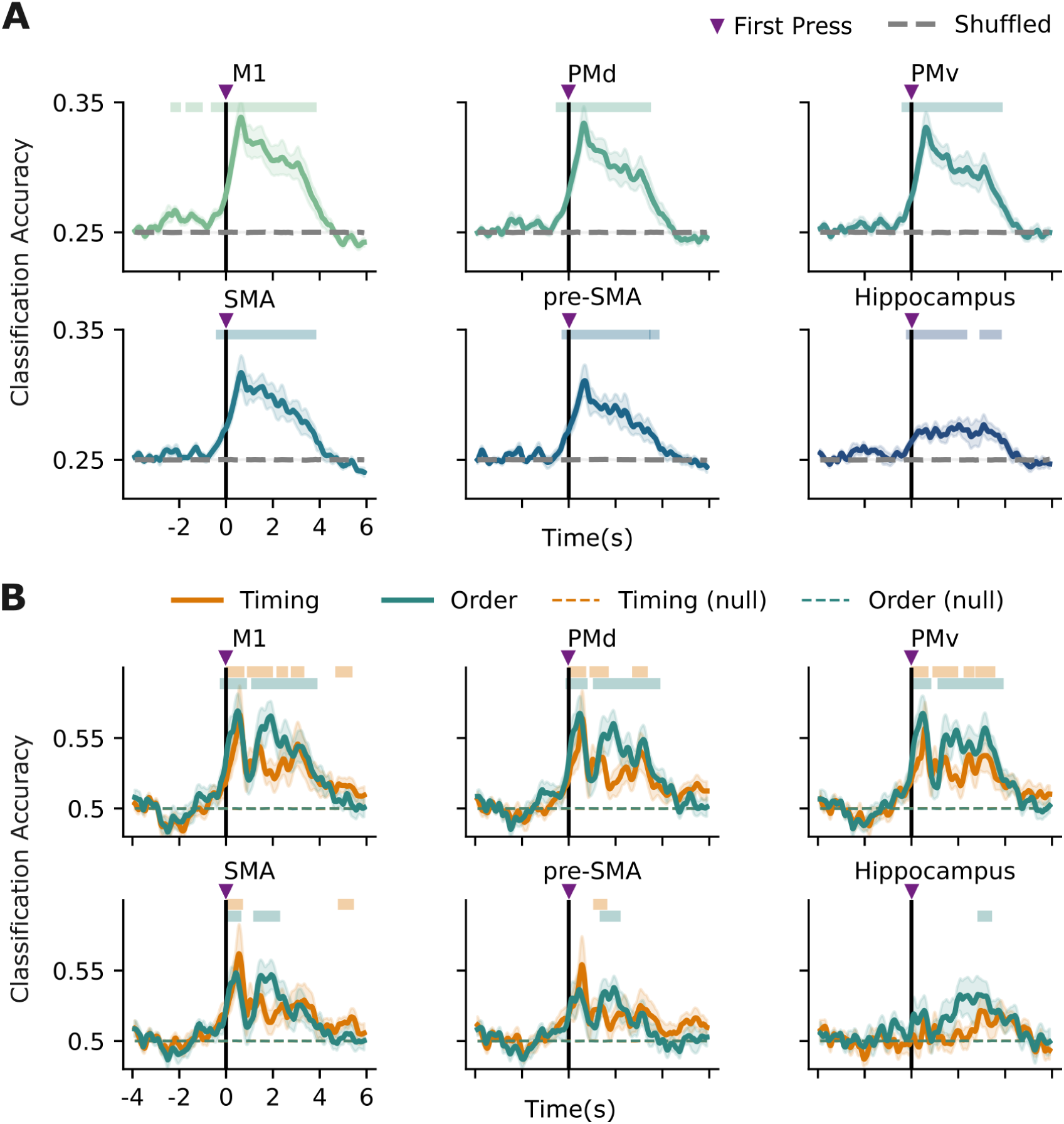
Movement-aligned sliding-window LDA decoding. (A) Sequence-identity decoding across regions. (B) Decoding of temporal-interval and finger-order factors. Top bars in both plots indicate time points at which decoding was significantly above the permutation-derived chance level, shown by the dashed line.

**Figure S4:**
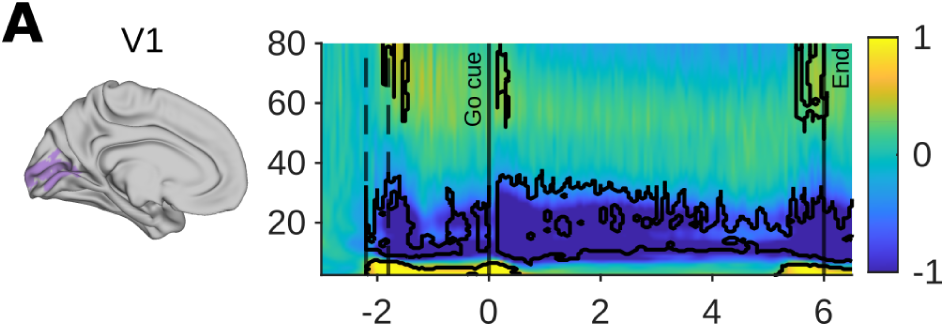
Time-frequency representation for V1. (A) Left: area mask used for source reconstruction. Right: TFR with significant clusters (relative to baseline) outlined.

**Figure S5:**
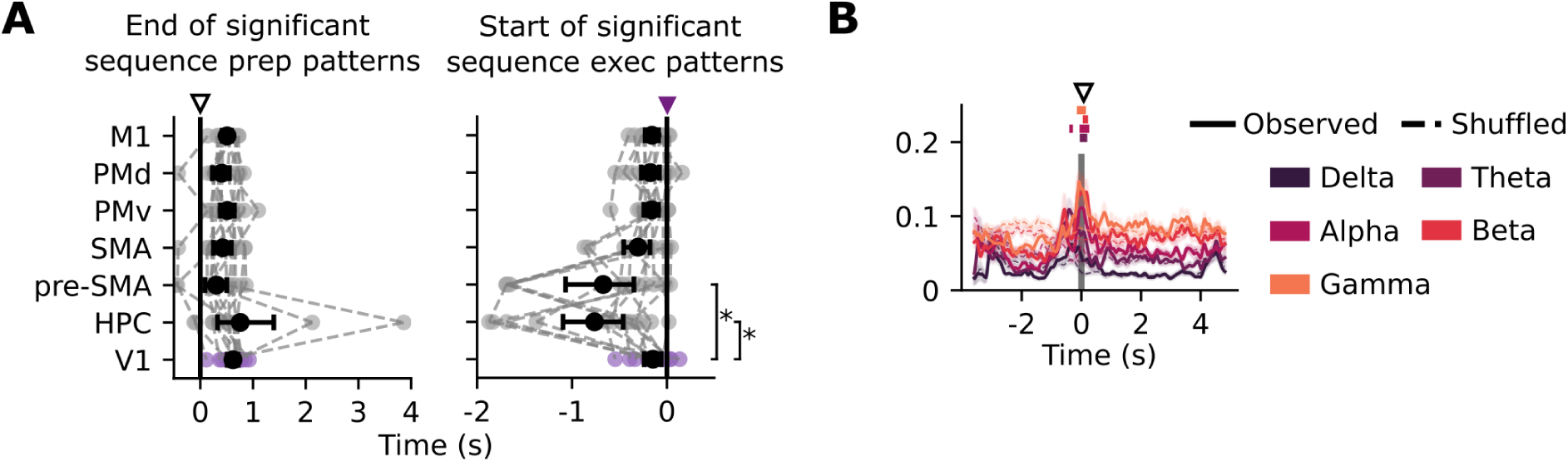
Phase-specific LDA decoding in V1. (A) Timing of the end of significant sequence-preparation decoding and the onset of significant sequence-execution decoding across regions. Purple indicates V1, whereas grey indicates the other regions. Brackets indicate significant differences between V1 and the other regions. (B) Regression analysis relating time–frequency activity to the movement-aligned LDA accuracy curve across frequency bands.

**Figure S6:**
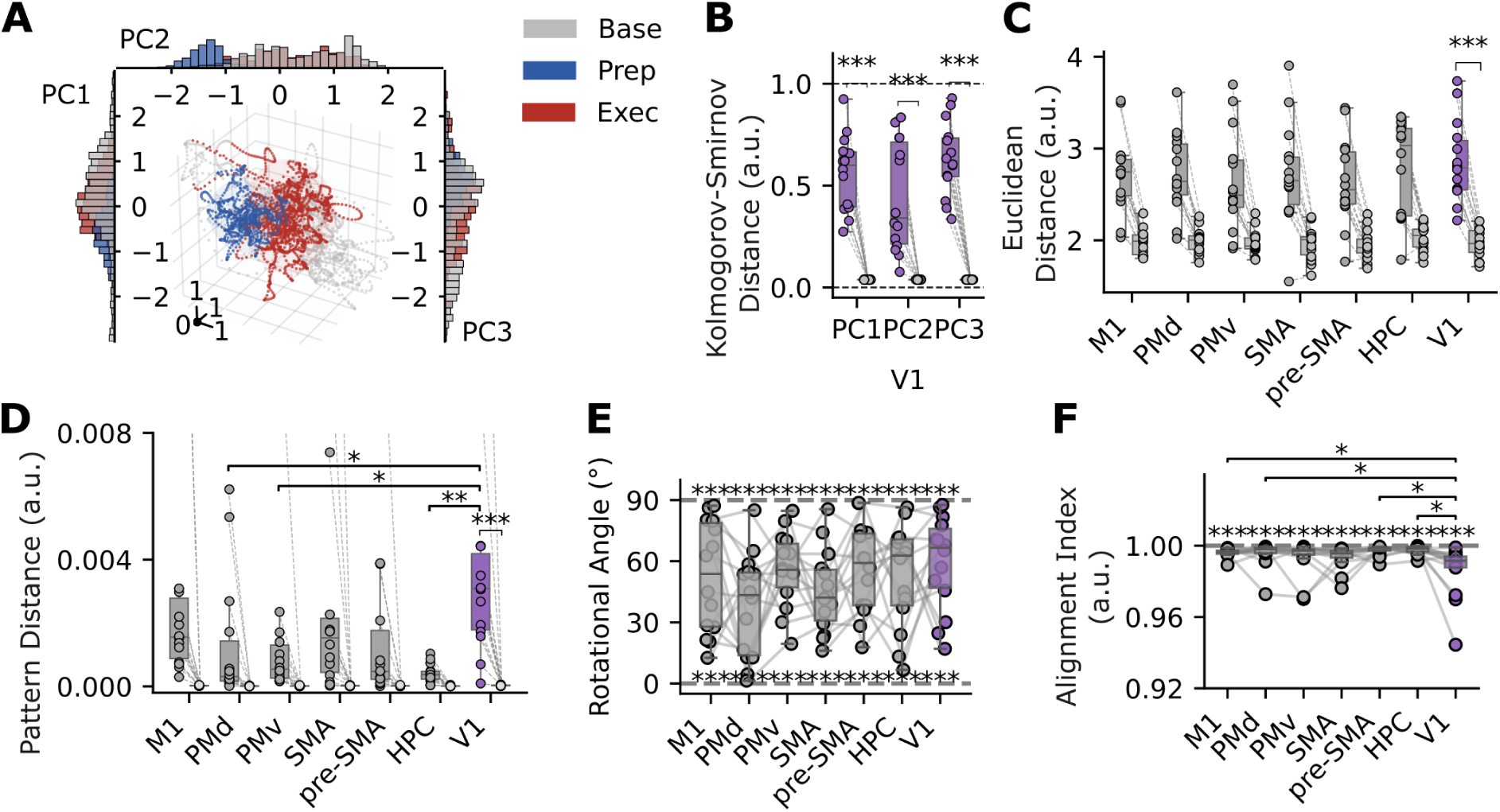
PCA analysis for V1. (A) Principal-component representation of V1 activity. (B) Kolmogorov–Smirnov distance between the preparation and execution distributions along each principal component. (C) Euclidean distance between preparatory- and execution-related activity across regions in PCA space. (D) Pattern distance between preparatory- and execution-related activity across regions. Brackets and asterisks denote significant pairwise differences in observed distance. (E) Rotational angle between maximum direction of variance for preparation and execution dynamics for V1. (F) Subspace-alignment index between preparatory- and execution-related activity.

**Figure S7:**
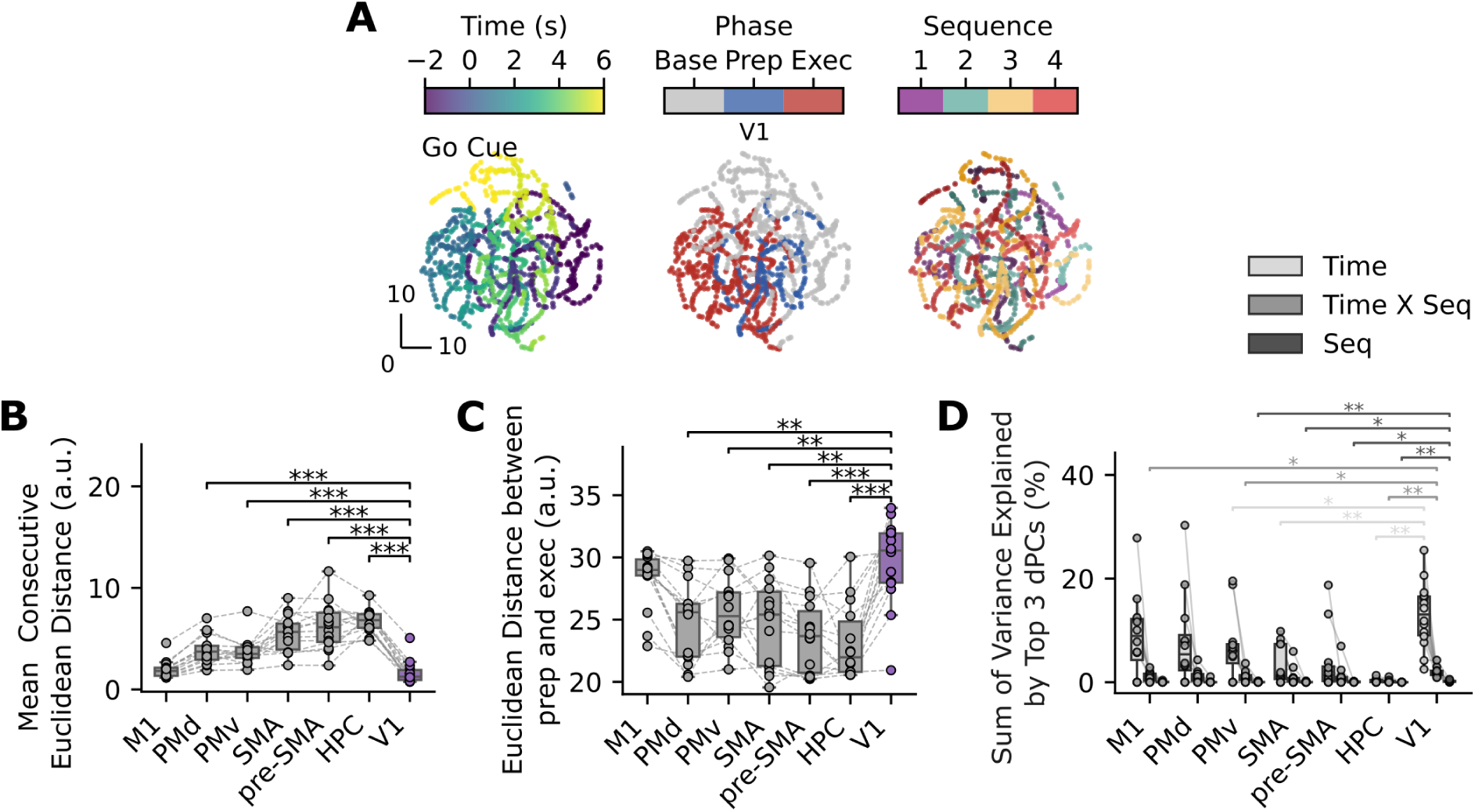
Covariance analysis for V1. (A) Three views of the same t-SNE embedding of V1 covariance matrices, with points coloured by continuous time (left), task phase (middle), or sequence identity (right). (B) Mean Euclidean distance between temporally consecutive points in the t-SNE embedding, averaged across sequences and participants. Brackets indicate significant pairwise differences between V1 and the other regions. (C) Euclidean distance between the preparation- and execution-phase clusters. (D) dPCA results showing the variance explained by each marginalisation. Brackets indicate significant differences between marginalisations.

**Figure S8:**
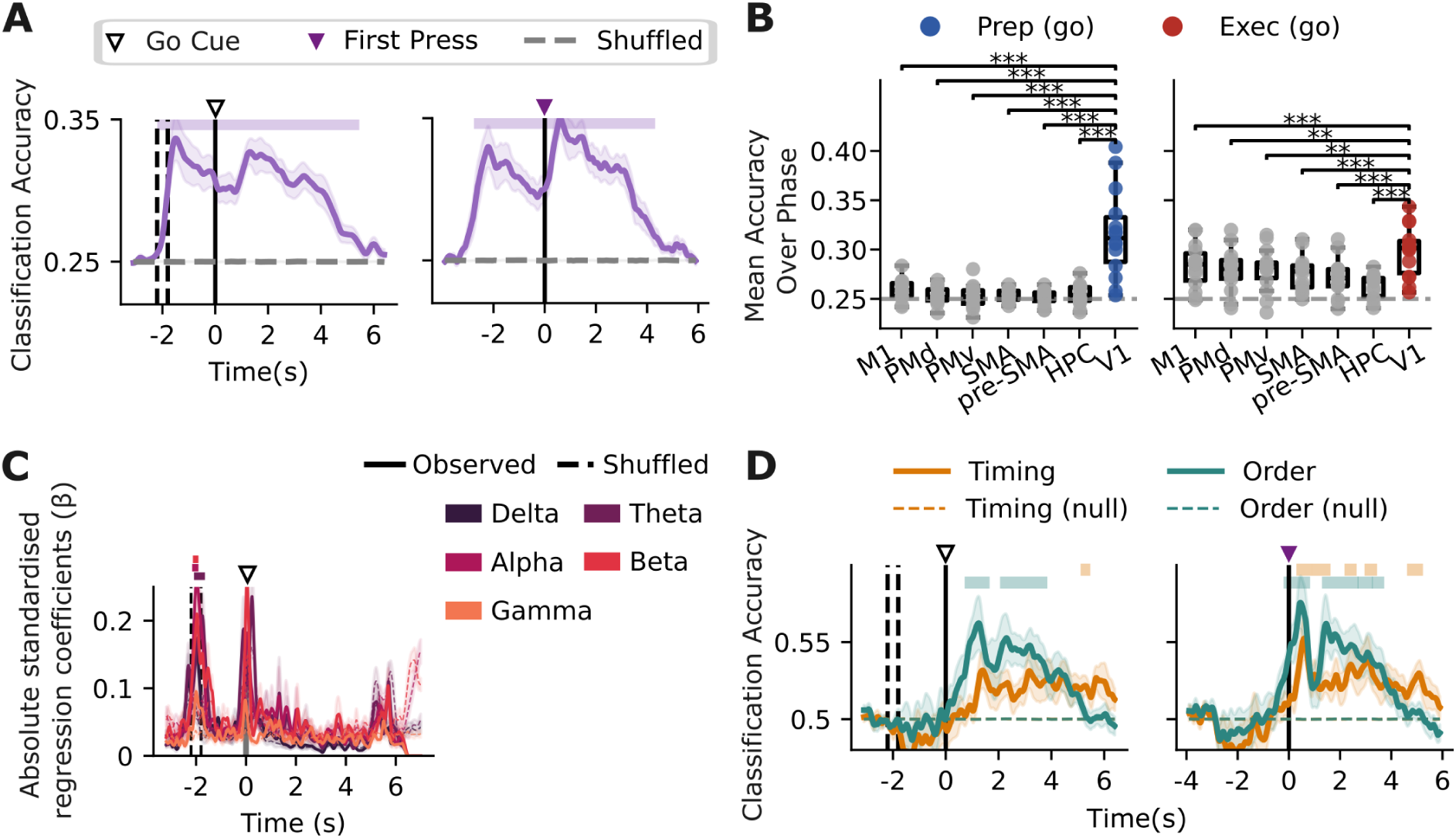
Sequence and factor decoding across regions and frequency bands for V1. (A) Sliding-window LDA decoding of sequence identity aligned to the Go Cue and movement onset. Top bars indicate significant decoding periods. (B) Mean sequence-decoding accuracy averaged across the preparation and execution phases. Colour indicates V1, whereas grey indicates motor regions and the hippocampus. Brackets indicate significant pairwise differences between V1 and other regions. (C) Association between time–frequency amplitude and sequence-decoding accuracy across frequency bands. Top bars indicate significant effects. (D) Sliding-window decoding of timing and finger-order factors, aligned to the Go Cue and movement onset.

**Table S1:** Significant time–frequency clusters by region compared to baseline for V1.

| Region | p | Mean dB change<br>to baseline | Frequency<br>range (Hz) | Time range<br>(s) | Peak/Trough<br>(time, Hz) | Peak dB |
| --- | --- | --- | --- | --- | --- | --- |
| V1 | < .001 | -1.27 | (6.0, 38.0) | (0.15, 6.8) | (6.0, 10) | -4.25 |
|  | < .001 | -1.15 | (8.5, 37.0) | (-2.2, -0.35) | (-1.5, 10.5) | -3.21 |
|  | .006 | -0.91 | (10.5, 32.5) | (-0.3, 0) | (-0.2, 12) | -1.54 |
|  | < .001 | 0.98 | (2.5, 7.5) | (-2.2, 0.55) | (0.1, 5) | 1.46 |
|  | .003 | 0.73 | (2.5, 6.0) | (5.1, 6.75) | (5.8, 3) | 1.04 |
|  | .008 | 0.30 | (48.0, 80.0) | (5.7, 6.05) | (5.9, 64) | 0.40 |
|  | .016 | 0.19 | (50.5, 80.0) | (5.5, 5.7) | (5.6, 70) | 0.30 |
|  | .018 | 0.29 | (52.0, 80.0) | (0.15, 0.35) | (0.15, 61) | 0.39 |
|  | .020 | 0.36 | (62.0, 80.0) | (-1.9, -1.7) | (-1.7, 80) | 0.53 |
|  | .021 | 0.37 | (54.0, 80.0) | (-1.55, -1.45) | (-1.5, 71) | 0.52 |
|  | .048 | 0.31 | (54.0, 80.0) | (5.75, 5.8) | (5.75, 69) | 0.38 |

**Table S2:** Significant clusters from regressions between movement-onset-aligned execution-state prediction curves and spectral power in V1 across canonical frequency bands.

| Area | Band | Time window 1 | Time window 2 |
| --- | --- | --- | --- |
| V1 | Theta | -0.12–0.13 ($p = .018$) | – |
| | Alpha | -0.47–-0.35 ($p = .046$) | -0.13–0.20 ($p = .0025$) |
| | Beta | -0.02–0.16 ($p = .029$) | – |
| | Gamma | -0.22–0.09 ($p = .010$) | – |

**Table S3:** Significant clusters from regressions between Go Cue-aligned sequence-decoding accuracy curves and spectral power in V1 across canonical frequency bands.

| Area | Band | Time window 1 | Time window 2 |
| --- | --- | --- | --- |
| V1 | Theta | -2.09–-1.65 ($p = .003$) | – |
| | Delta | -2.11–-1.90 ($p = .011$) | – |
| | Alpha | -2.16–-1.91 ($p = .008$) | -1.83–-1.71 ($p = .039$) |
| | Beta | -2.13–-1.91 ($p = .009$) | -1.82–-1.69 ($p = .026$) |
| | Gamma | -2.03–-1.92 ($p = .021$) | – |

